# The *Legionella* autoinducer LAI-1 is delivered by outer membrane vesicles to promote inter-bacterial and inter-kingdom signaling

**DOI:** 10.1101/2023.08.22.554324

**Authors:** Mingzhen Fan, Patrick Kiefer, Paul Charki, Christian Hedberg, Jürgen Seibel, Julia A. Vorholt, Hubert Hilbi

## Abstract

*Legionella pneumophila* is an environmental bacterium, which replicates in amoeba but also in macrophages, and causes a life-threatening pneumonia called Legionnaires’ disease. The opportunistic pathogen employs the α-hydroxyketone compound LAI-1 (*Legionella* autoinducer-1) for intra-species and inter-kingdom signaling. LAI-1 is produced by the autoinducer synthase LqsA, but it is not known, how LAI-1 is released by the pathogen. Here, we use a *V. cholerae* luminescence reporter strain and liquid chromatography-tandem mass spectrometry (LC-MS/MS) to detect bacteria-produced and synthetic LAI-1. Ectopic production of LqsA in *E. coli* generated LAI-1, which partitions to outer membrane vesicles (OMVs), and slightly reduces OMV size. These *E. coli* OMVs trigger luminescence of the *V. cholerae* reporter strain and inhibit the migration of *Dictyostelium discoideum* amoeba. Overexpression of *lqsA* in *L. pneumophila* under the control of strong stationary phase promoters (P*_flaA_* or P*_6SRNA_*), but not under control of its endogenous promoter (P*_lqsA_*), produces LAI-1, which is detected in purified OMVs. These *L. pneumophila* OMVs trigger luminescence of the *Vibrio* reporter strain and inhibit *D. discoideum* migration. *L. pneumophila* OMVs are smaller upon overexpression of *lqsA* or upon addition of LAI-1 to growing bacteria, and therefore, LqsA affects OMV production. The overexpression of *lqsA* but not a catalytically inactive mutant promotes intracellular replication of *L. pneumophila* in macrophages, indicating that intracellularly produced LA1-1 modulates the interaction in favour of the pathogen. Taken together, we provide evidence that *L. pneumophila* LAI-1 is secreted through OMVs and promotes inter-bacterial communication as well as interactions with eukaryotic host cells.

**Originality - Significance Statement:** Inter-kingdom signaling involving low molecular weight bacterial compounds that are detected by eukaryotic cells represents an important, yet incompletely understood aspect of pathogen-host interactions. In many cases, the small signaling molecules are produced in only little amounts, their secretion mechanism is not known, and their effects on eukaryotic host cells are barely studied. Here, we reveal that the α-hydroxyketone compound LAI-1 of *L. pneumophila* is released from the bacteria by outer membrane vesicles, which promote inter-bacterial communication as well as inter-kingdom signaling.

## Introduction

The gram-negative bacterium *Legionella pneumophila* is an opportunistic pathogen, which upon inhalation of bacteria-laden aerosols replicates in lung macrophages and causes a life-threatening pneumonia termed Legionnaires’ disease (1–3). In the environment, *L. pneumophila* persists and replicates in free-living protozoa (4–6). Intriguingly, *L. pneumophila* subverts the bactericidal potential of macrophages and amoeba in a similar manner and within these evolutionary distant phagocytes forms an ER-associated replication-permissive compartment called the *Legionella*-containing vacuole (LCV) (7–10). To govern LCV formation, *L. pneumophila* employs the Icm/Dot type IV secretion system (T4SS), which translocates more than 300 “effector” proteins into eukaryotic host cells, where they subvert various processes, including trafficking pathways, cytoskeleton dynamics, signal transduction, and metabolism (9,11–16).

*L. pneumophila* employs the *Legionella* quorum sensing (Lqs) system for small molecule intra-species and inter-kingdom signaling (**Fig. 1**) (17,18). The quorum sensing system is rather broadly distributed among environmental bacteria, including the families Legionellaceae, Vibrionaceae, Burkholderiaceae, Chlorobiaceae, and Oxalobacteraceae (17,19), and fairly conserved among *Legionella* spp.: 19 out of 58 species harbor a complete *lqs* cluster (20).

**Figure 1.**
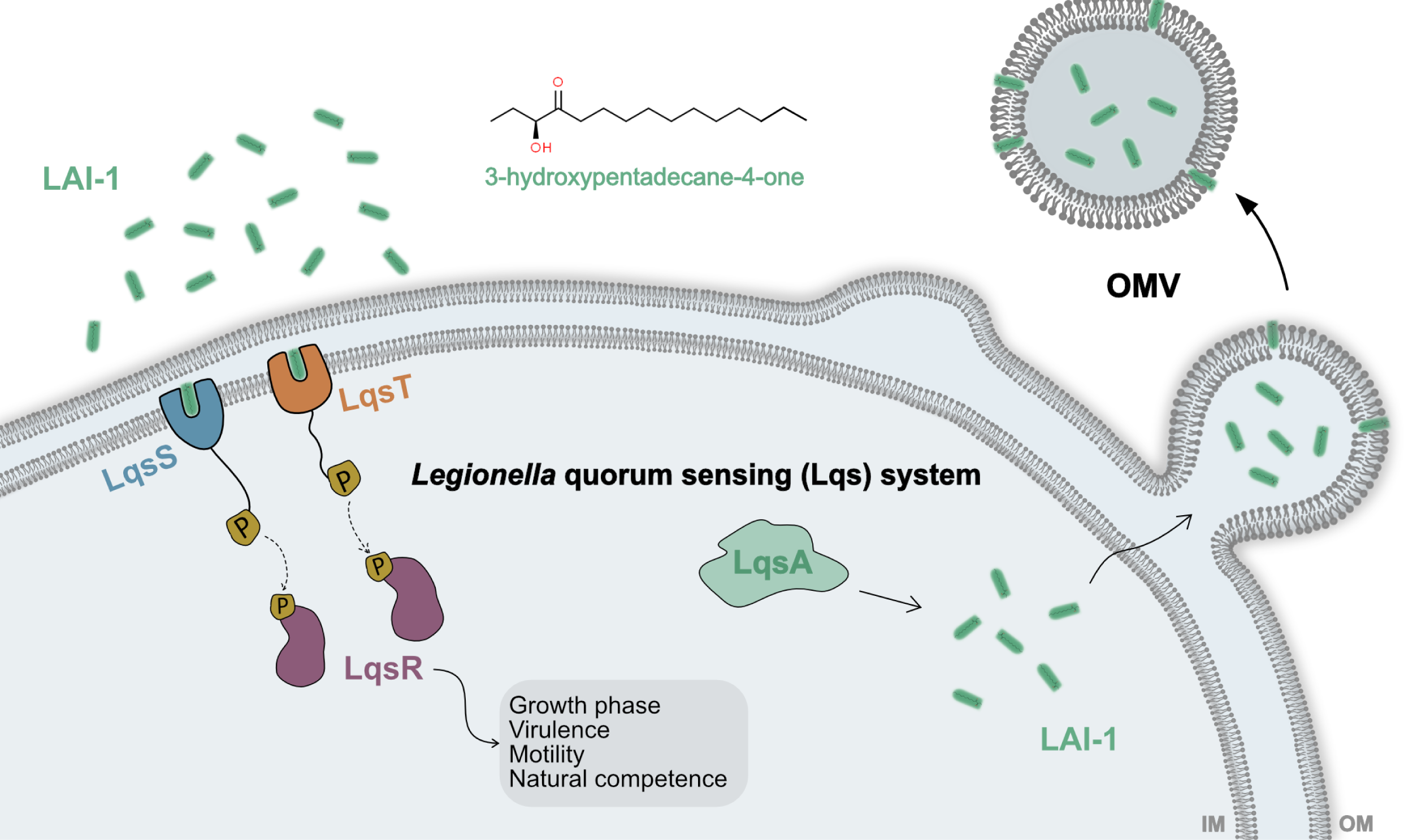
The *L. pneumophila* Lqs system: Production, detection, and release of LAI-1. The Lqs (*Legionella* quorum sensing) system produces, detects, and responds to the small signaling molecule LAI-1 (*Legionella* autoinducer-1, 3-hydroxypentadecane-4-one). The system comprises the autoinducer synthase LqsA, the cognate membrane-bound sensor kinases LqsS and LqsT, and the prototypic response regulator LqsR. LAI-1 produced by LqsA partitions into outer membrane vesicles (OMVs), through which the hydrophobic compound is released from the bacteria to promote inter-bacterial as well as inter-kingdom signaling.

The Lqs system includes the pyridoxal-5′-phosphate (PLP)-dependent autoinducer synthase LqsA (21), which is 41% identical to *V. cholerae* CqsA (22,23). Furthermore, the system comprises the homologous sensor histidine kinases LqsS (24) and LqsT (25), and the cognate response regulator LqsR (26,27), which dimerizes upon phosphorylation (28,29). LqsS negatively regulates a pleiotropic transcription factor termed *Legionella* virulence and biofilm regulator (LvbR), which controls *L. pneumophila* virulence, biofilm architecture and natural competence for DNA uptake (30). LvbR also regulates the nitric oxide (NO) sensor and di-guanylate cyclase inhibitor Hnox1, and thus, positively regulates the production of the second messenger cyclic di-guanosine monophosphate (c-di-GMP). Accordingly, quorum sensing is linked to c-di-GMP signaling in *L. pneumophila* (30,31).

The Lqs system produces, detects, and responds to the low molecular weight compound LAI-1 (*Legionella* autoinducer-1, 3-hydroxypentadecane-4-one) (17,32) (**Fig. 1**). The quorum sensing system and synthetic LAI-1 regulate various traits of *L. pneumophila*, such as motility and flagellum production (33), virulence (18,34), the bacterial growth phase switch and temperature-dependent culture density (26,35), expression of a “fitness island”, and natural competence for DNA uptake (18,34). Moreover, the Lqs-LvbR signaling network regulates the migration of *Acanthamoeba castellanii* amoeba through *L. pneumophila* biofilms (36), as well as – upon exposure of *Dictyostelium discoideum* amoeba, macrophages, or epithelial cells to *L. pneumophila* – the motility of eukaryotic cells (37). LAI-1 is rather hydrophobic and might partition into the aqueous or lipid bacterial compartments (21,33). It is not known how LAI-1 is secreted and delivered by *L. pneumophila*.

Gram-negative bacteria form and release so-called outer membrane vesicles (OMVs), which are spherical vesicles of ca. 10-300 nm diameter confined by a membrane bilayer with lipopolysaccharide (LPS) in the outer leaflet (38,39). OMVs are produced and released by planktonic, biofilm, and intracellular bacteria under different physiological conditions, and they transport proteins, hydrophobic small molecules, and nucleic acids between bacteria or between bacteria and eukaryotic cells (40,41). OMVs adopt a plethora of functions, including disposal of “waste” material (proteins, lipids, peptidoglycan), secretion of virulence factors (toxins, proteases, lipases), nutrient scavenging (carbon sources, micronutrients: e.g., iron), inactivation of antibiotics, titration of phages, transport of signaling molecules, as well as delivery of DNA and regulatory RNA (42). While the production of OMVs comes with many advantages for the producing bacteria, it also comes with a price, as OMVs are complex and energetically costly produced, and they elicit possibly adverse immune responses (43).

*L. pneumophila* produces OMVs, which are enriched in virulence-relevant proteins such as proteolytic and lipolytic enzymes, the macrophage infectivity potentiator (Mip) protein, Icm/Dot components and substrates, as well as the major flagellum component flagellin (44,45). These OMVs can fuse with eukaryotic membranes, but do not kill host cells and rather promote the growth of amoeba, activate mammalian cells, and modulate their cytokine response (44,46–48). The host cell’s innate immune response is also targeted by small RNAs delivered by *L. pneumophila* OMVs (49). Moreover, *L. pneumophila* OMVs bind to and are internalized by macrophages (46,50), where – independently of the Icm/Dot T4SS – they inhibit phagosome-lysosome fusion (51) and promote intracellular bacterial replication at later time points of infection (47).

In this study, we tested the hypothesis that the *L. pneumophila* signaling compound LAI-1 partitions to OMVs and is secreted by these vesicles. We provide evidence that LAI-1 partitions to and is secreted by OMVs formed in either *E. coli* or *L. pneumophila* overexpressing the autoinducer synthase gene *lqsA*. These OMVs mediate intra-bacterial as well as inter-kingdom communication, and overexpression of *lqsA* but not a catalytically inactive form promotes intracellular replication of *L. pneumophila* in macrophages.

## Results

### LAI-1 detection by a Vibrio reporter strain and LC-MS/MS

The *L. pneumophila* signaling compound LAI-1 has been identified as 3-hydroxypentadecane-4-one by liquid chromatography-tandem mass spectrometry (LC-MS/MS) upon overexpression of *lqsA* controlled by the P*_tac_* promoter in *E. coli* and *L. pneumophila* (21). Under the conditions tested, *L. pneumophila* strains did not produce endogenous LAI-1 detectable by MS or by the *V. cholerae* α-hydroxyketone luminescence reporter strain MM920 (21) (**Fig. 2A**). Upon P*_tac_*-controlled overexpression of *lqsA* in *E. coli*, *L. micdadei* or *L. pneumophila* growing on CYE agar plates, the *V. cholerae* strain MM920 detected a signal produced by *E. coli* and *L. micdadei*, but not by several *L. pneumophila* strains (JR32, Corby, clinical isolate #883) (**Fig. 2A**). However, upon growing in AYE broth, *L. pneumophila* overexpressing *lqsA* produced a signal, which was detected by the *V. cholerae* reporter strain in culture supernatants, albeit only during a short period of time during growth (**Fig. S1**). Contrarily, the signal produced by *L. pneumophila* overexpressing *cqsA* was robustly detected by the *V. cholerae* reporter strain, regardless of whether *L. pneumophila* was grown on agar plates or in broth (**Fig. 2A, Fig. S1**).

**Figure 2.**
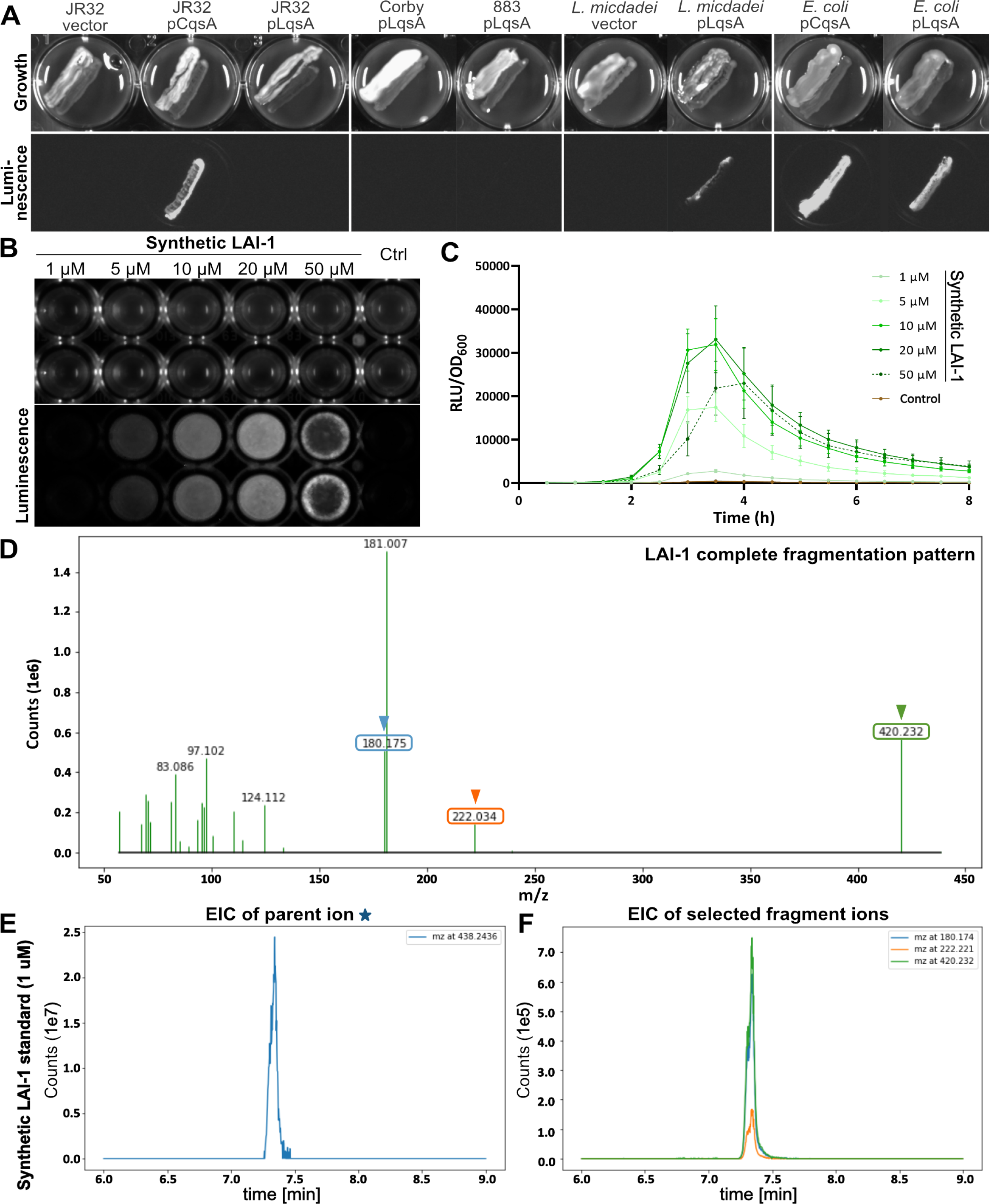
LAI-1 detection by *V. cholerae* MM920 and LC-MS/MS. (**A**-**C**) Luminescence of the *V. cholerae* α-hydroxyketone reporter strain MM920. (**A**) *L. pneumophila* strains (JR32, Corby, clinical isolate 883), *L. micdadei*, or *E. coli* harboring an empty vector (pTS-10) or plasmids expressing *lqsA* (pTS-2) or *cqsA* (pTS-6) under control of the P*_tac_* promoter were grown on CYE agar in 24 well plates for 2 days, before the *V. cholerae* reporter strain MM920 was streaked out in parallel. After another day, growth was assessed (upper panels), and bioluminescence was determined with a luminometer (lower panels). *V. cholerae* MM920 was treated with the concentrations of synthetic LAI-1 indicated, DMSO was included as negative control, and luminescence intensity was (**B**) visualized by a gel documentation system (15 min exposure time), or (**C**) measured by a plate reader (30°C, 8 h). RLU, relative light units. (**D**-**F**) MS fragment spectra of LAI-1. (**D**) The fragment ions at m/z 180.174 and 222.221 cover the complete structure of LAI-1 and exhibit high specificity. For detection of LAI-1, the focus was placed on fragment ions at m/z 180.174 (light blue arrowhead), 222.221 (orange arrowhead), and 420.232 (green arrowhead). Positive detection and identification results were obtained from the extracted ion chromatogram (EIC) of (**E**) the parent ion at m/z 438.24, along with (**F**) compound-specific fragment ions at m/z 180.174 (light blue line), 222.221 (orange line), and 420.232 (green line). LC-MS/MS analysis was performed with 1 µM synthetic LAI-1 as a standard.

Given the apparently low amounts of LAI-1 produced by *L. pneumophila*, we next tested whether and with which sensitivity the *V. cholerae* reporter strain detects LAI-1. To this end, 1-50 µM synthetic LAI-1 was added to the reporter strain and luminescence was measured (**Fig. 2B-C**). 1-20 µM synthetic LAI-1 triggered the luminescence response by the *V. cholerae* reporter strain in a dose-dependent manner, indicating that the reporter strain indeed recognizes the *L. pneumophila* signaling molecule. 50 µM LAI-1 interfered with the growth of *V. cholerae*, and accordingly, the luminescence emitted by the reporter strain was reduced. The detection limit for LAI-1 of the *V. cholerae* luminescence reporter assay was ca. 1 µM (**Fig. 2C**).

Synthetic LAI-1 was also detected by LC-MS/MS (**Fig. 2D-F, Fig. S2**). After oximation with O-(2,3,4,5,6-pentafluorobenzyl) hydroxylamine hydrochloride (O-PFB), a standard of 1 µM LAI-1 was analyzed. LAI-1 was detected and identified by the extracted ion chromatogram, showing the parent ion at m/z 438.24 (**Fig. 2E, Fig. S2**), along with compound-specific fragment ions at m/z 180.174, 222.221, and 420.232 (**Fig. 2F, Fig. S2**). The detection limit of the LC-MS/MS approach for LAI-1 was ca. 10 fmol corresponding to 1µl of a 10 nM standard solution.

### LAI-1 partitions to OMVs of E. coli ectopically expressing lqsA

To assess whether LAI-1 affects OMV formation and/or partitions into OMVs, we ectopically expressed *lqsA* in *E. coli* under the control of the P*_tac_* promoter. OMVs were isolated from bacterial culture supernatants and enriched by ultracentrifugation as outlined in the Experimental Procedures section (**Fig. 3A**). Compared to the control strain, *E. coli* expressing *lqsA* produced a population of OMVs, the majority of which ranging from approximately 10-60 nm in diameter, with slightly reduced average size (**Fig. 3B**). Hence, the ectopic production of LqsA in *E. coli* seems to affect the formation of OMVs.

**Figure 3.**
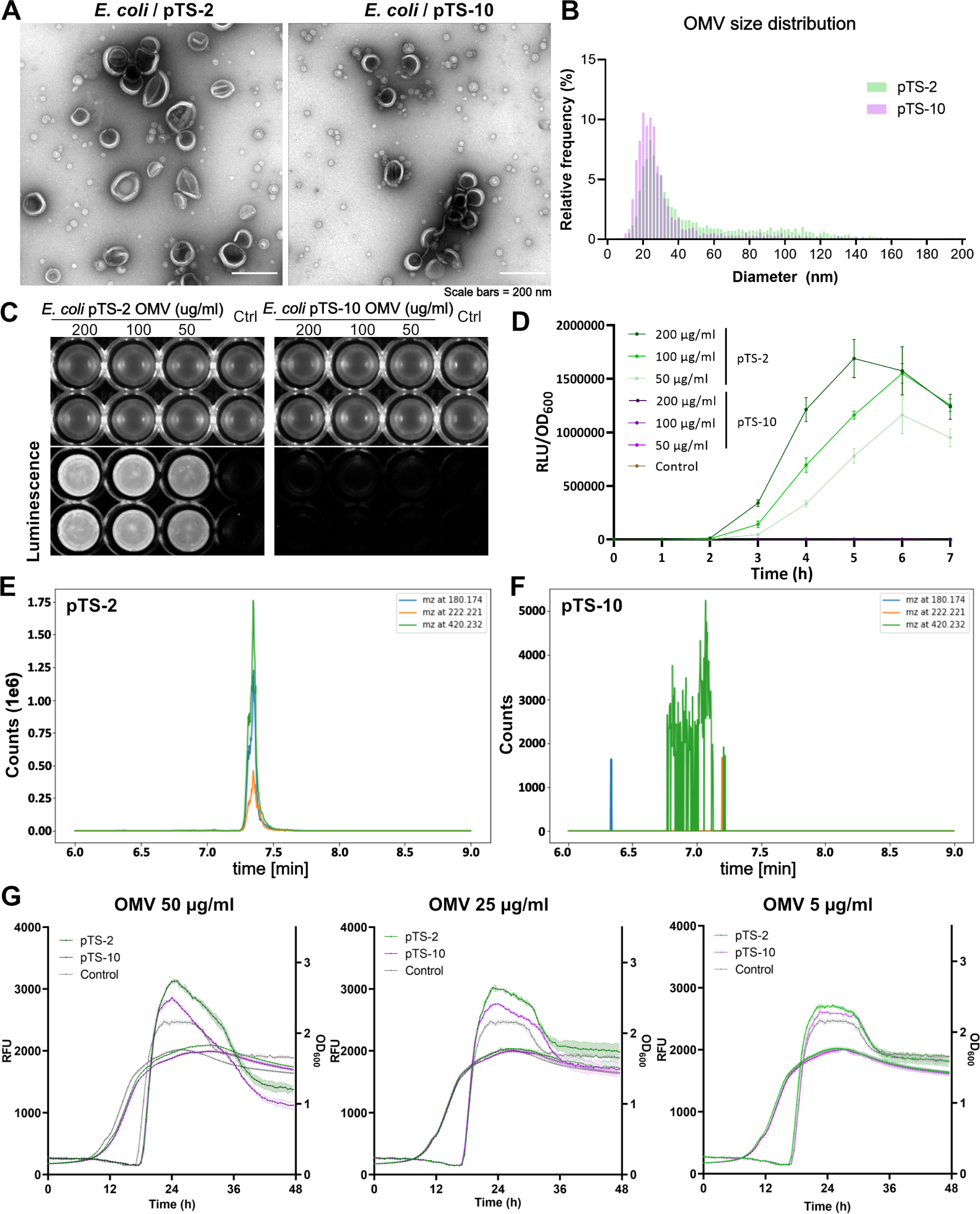
LAI-1 partitions to OMVs of *E. coli* ectopically expressing *lqsA* and promotes inter-bacterial signaling. OMVs of *E. coli* TOP10 harboring pTS-2 (P_tac_-*lqsA*) or pTS-10 (empty vector) were isolated from overnight cultures induced with 1 mM IPTG for 4 h. **(A)** Negative staining transmission electron microscopy (TEM) images of OMVs derived from *E. coli* harboring pTS-2 (left) or pTS-10 (right). (**B**) Size distribution analysis of OMVs of *E. coli* pTS-2 or pTS-10. **(C-D)** Luminescence of *V. cholerae* MM920 mixed with OMVs from *E. coli* harboring pTS-2 or pTS-10 of the protein concentrations indicated (control: DMSO). Intensity was (**C**) visualized by a gel documentation system (15 min exposure time), or (**D**) measured by a plate reader (30°C, 7 h). RLU, relative light units. **(E-F**) LC-MS/MS analysis of OMVs from *E. coli* pTS-2 or pTS-10 (1 mg protein). Extracted ion chromatograms (EICs) of fragment ions at m/z 180.174 (light blue), 222.221 (orange), and 420.232 (green) indicated the presence of LAI-1 in OMVs of *E. coli* pTS-2 (P*_tac_-lqsA*) but not in OMVs of *E. coli* pTS-10. (**G**) *L. pneumophila* expressing P*_flaA_*-*gfp* was treated with OMVs of *E. coli* harboring pTS-2 (P*_tac_-lqsA*) or pTS-10 (empty vector). OMVs from *E. coli* pTS-2 induced the expression of *gfp* under the control of P*_flaA_*in a dose-dependent manner.

We then exposed the *V. cholerae* reporter strain MM920 to OMV preparations purified from *E. coli* expressing *lqsA* or not (**Fig. 3C**). OMV preparations (50-200 µg protein/ml) purified from *E. coli* expressing *lqsA* resulted in a dose-dependent luminescence signal, while OMV samples prepared from the *E. coli* control strain or untreated reporter strain did not produce any luminescence (**Fig. 3D**). LAI-1 was also detected by LC-MS/MS in OMV preparations from *E. coli* expressing *lqsA* (**Fig. 3E**) but not from the *E. coli* control strain (**Fig. 3F**, **Table 1**).

**Table 1.**
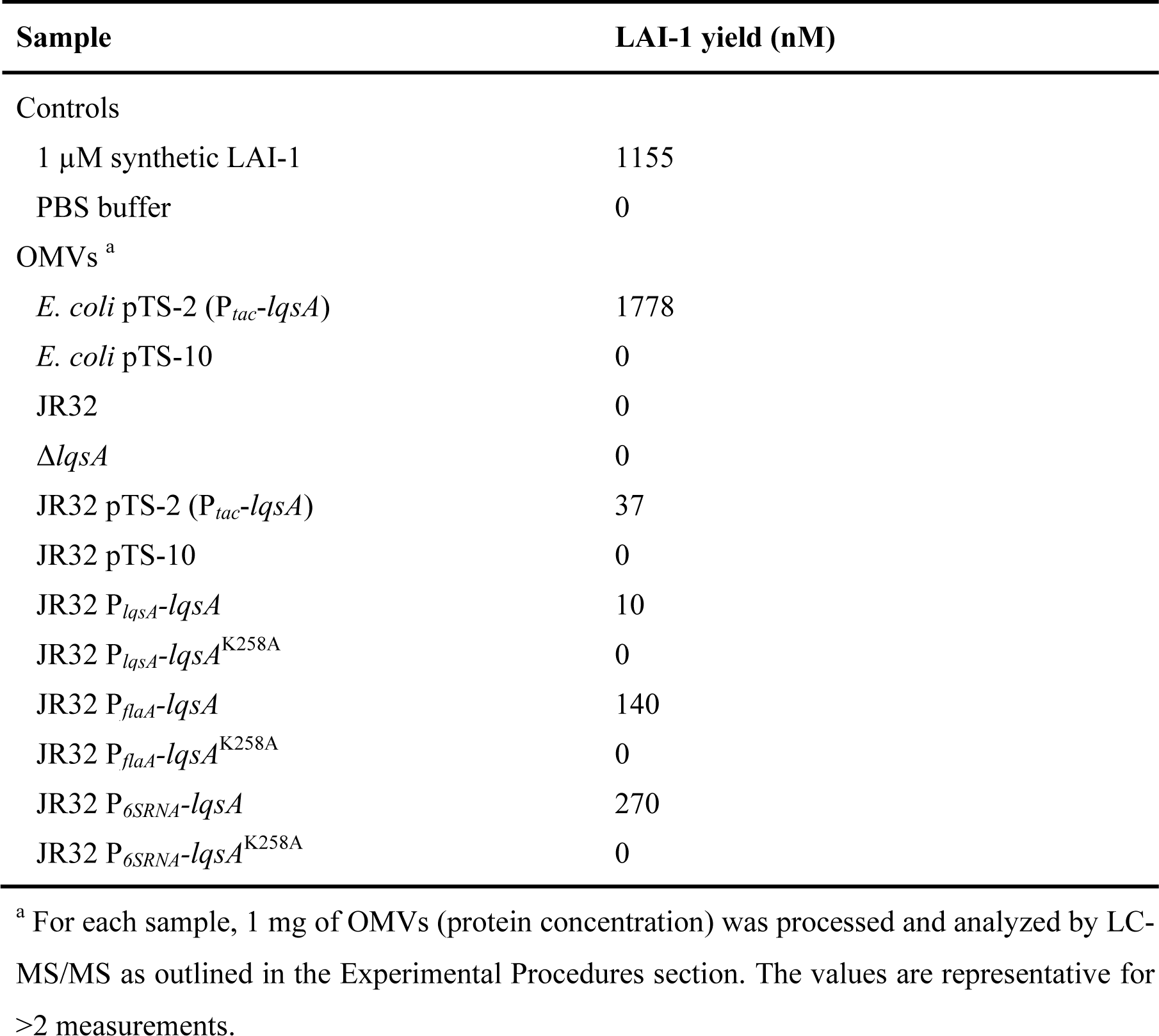
LAI-1 detection of OMV samples by LC-MS/MS.

| Sample | LAI-1 yield (nM) |
| --- | --- |
| Controls |  |
| 1 μM synthetic LAI-1 | 1155 |
| PBS buffer | 0 |
| OMVs <sup>a</sup> |  |
| *E. coli* pTS-2 (*P<sub>tac</sub>-lqsA*) | 1778 |
| *E. coli* pTS-10 | 0 |
| JR32 | 0 |
| $\Delta lqsA$ | 0 |
| JR32 pTS-2 (*P<sub>tac</sub>-lqsA*) | 37 |
| JR32 pTS-10 | 0 |
| JR32 *P<sub>lqsA</sub>-lqsA* | 10 |
| JR32 *P<sub>lqsA</sub>-lqsA*<sup>K258A</sup> | 0 |
| JR32 *P<sub>flaA</sub>-lqsA* | 140 |
| JR32 *P<sub>flaA</sub>-lqsA*<sup>K258A</sup> | 0 |
| JR32 *P<sub>6SRNA</sub>-lqsA* | 270 |
| JR32 *P<sub>6SRNA</sub>-lqsA*<sup>K258A</sup> | 0 |

<sup>a</sup> For each sample, 1 mg of OMVs (protein concentration) was processed and analyzed by LC- MS/MS as outlined in the Experimental Procedures section. The values are representative for >2 measurements.

OMV preparations purified from *E. coli* expressing *lqsA* also induced the expression of *gfp* under the control of P*_flaA_* in *L. pneumophila* in a dose-dependent manner (**Fig. 3G**). Using this readout, OMV samples prepared from the *E. coli* control strain barely induced the expression of P*_flaA_*-*gfp*. Accordingly, not only the *V. cholerae* reporter strain, but also *L. pneumophila* detects and responds to OMV-associated LAI-1. Taken together, the ectopic production of LqsA in *E. coli* produces LAI-1, which partitions to OMVs, affects OMV formation of the donor strain, and promotes inter-bacterial communication.

### E. coli OMVs containing LAI-1 inhibit the migration of D. discoideum

The migration of *D. discoideum* is reduced by pretreatment with 10 µM synthetic LAI-1 (37) (**Fig S3**). Based on this finding, we asked the question, whether *E. coli* OMVs containing LAI-1 would exhibit a comparable effect and promote inter-kingdom signaling. To this end, we treated *D. discoideum* amoeba with OMVs purified from *E. coli* expressing *lqsA* or not (**Fig. 4**). Compared to the MB medium control (**Fig. 4A**), the migration of the amoeba was inhibited in a dose-dependent manner by OMVs purified from *E. coli* expressing *lqsA* (**Fig. 4B**), but not by OMVs from *E. coli* harboring an empty plasmid (**Fig. 4C**). At the highest OMV concentration used (500 µg/ml protein), the velocity of the amoeba was reduced ca. 3-fold compared to the controls (**Fig. 4D**). Accordingly, OMV-associated LAI-1 not only promotes inter-bacterial communication but also inter-kingdom signalining.

**Figure 4.**
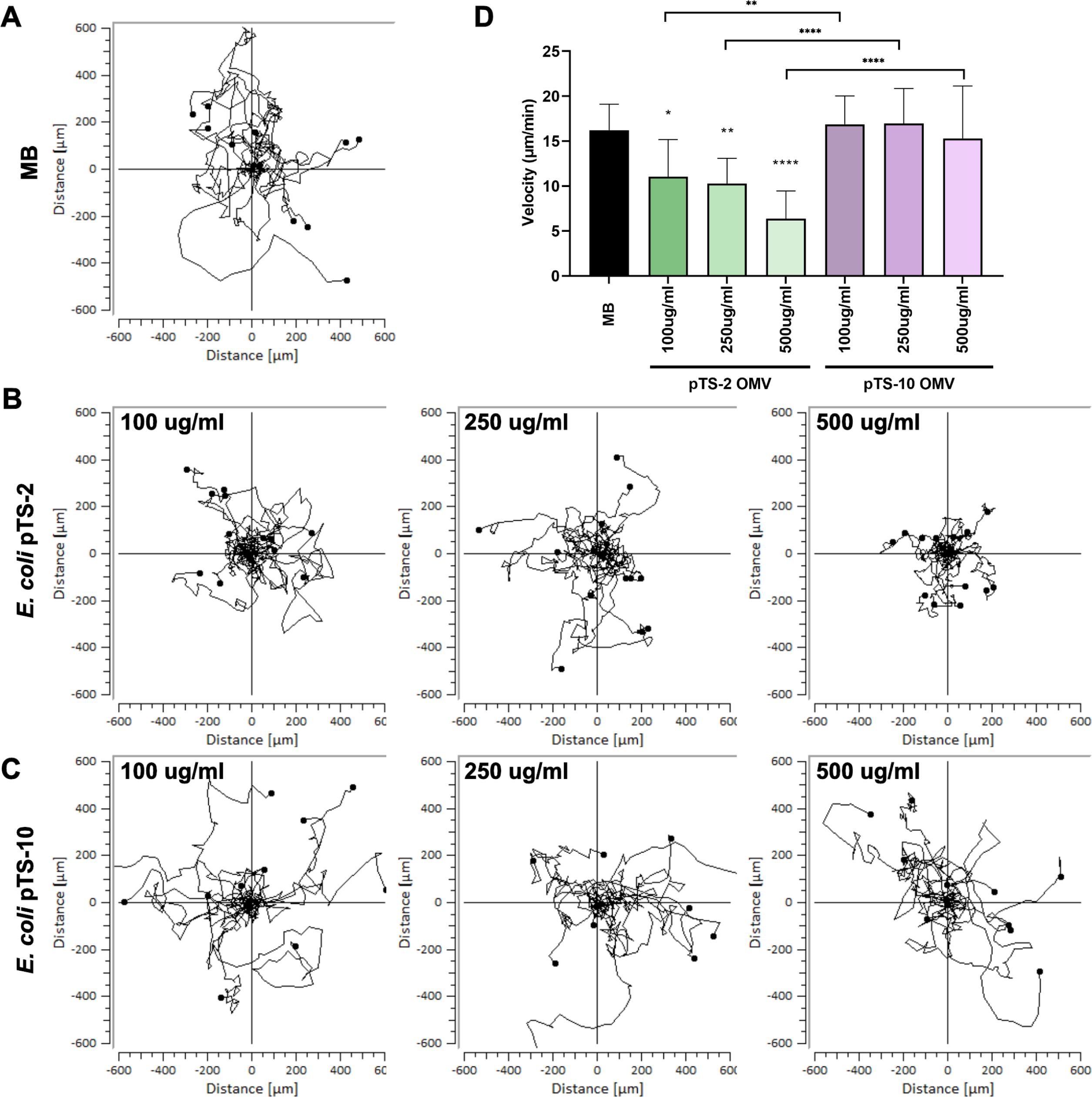
*E. coli* OMVs harboring LAI-1 inhibit the migration of *Dictyostelium.* *D. discoideum* amoeba were treated with OMVs of *E. coli* TOP10 harboring pTS-2 (P*_tac_*-*lqsA*) or pTS-10 (empty vector) with a final protein concentration of 100 µg/ml, 250 µg/ml, or 500 µg/ml. For each sample, the migration of 10-15 amoeba was tracked over 2 h and the velocity was quantified. Migration trajectories of amoeba treated with (**A**) MB medium (control), (**B**) OMVs of *E. coli* pTS-2, or (**C**) *E. coli* pTS-10. (**D**) Median of amoeba migration velocity. The data shown are velocity median of 10-15 amoeba per sample (*, *p* ≤ 0.1; **, *p* ≤ 0.01; ****, *p* ≤ 0.0001) and representative of 3 independent experiments.

### L pneumophila OMV production is regulated by LqsA and LAI-1

Similar to *E. coli* (**Fig. 3B**), the production of OMVs by *L. pneumophila* might be affected by LAI-1. In order to maximize the potential effect of LqsA and LAI-1 production on OMV formation, we thought to correlate P*_lqsA_* activity with OMV formation. In *L. pneumophila*, *lqsA* (25) and P*_lqsA_* (**Fig. S4**) are expressed from early stationary throughout later stationary growth phase, and therefore, we routinely isolated OMVs from *L. pneumophila* in stationary growth phase.

To test whether LqsA affects OMV formation of *L. pneumophila*, we first compared the OMV populations produced by the parental *L. pneumophila* strain JR32 or the Δ*lqsA* mutant strain (**Fig. 5A**). The strains JR32 and Δ*lqsA* produced similar OMV populations, with individual OMV ranging in size from approximately 20 to 120 nm in diameter and a median diameter of 62 nm (**Fig. 5B**). Hence, under the conditions used, the absence of LqsA does not seem to affect the formation of OMVs in *L. pneumophila*. We then exposed the *V. cholerae* reporter strain MM920 to OMV preparations purified from *L. pneumophila* JR32 or Δ*lqsA* (**Fig. 5C**). Strikingly, none of the OMV preparations tested at a concentration of 50-200 µg/ml produced a luminescence signal above the background level of the untreated reporter strain (**Fig. 5D**). Similarly, LAI-1 was also not detected by LC-MS/MS in OMV preparations from strains JR32 (**Fig. 5E**) or Δ*lqsA* (**Fig. 5F**, **Table 1**). Taken together, under the conditions tested, *L. pneumophila* does not produce and purified OMVs do not contain detectable levels of LAI-1.

**Figure 5.**
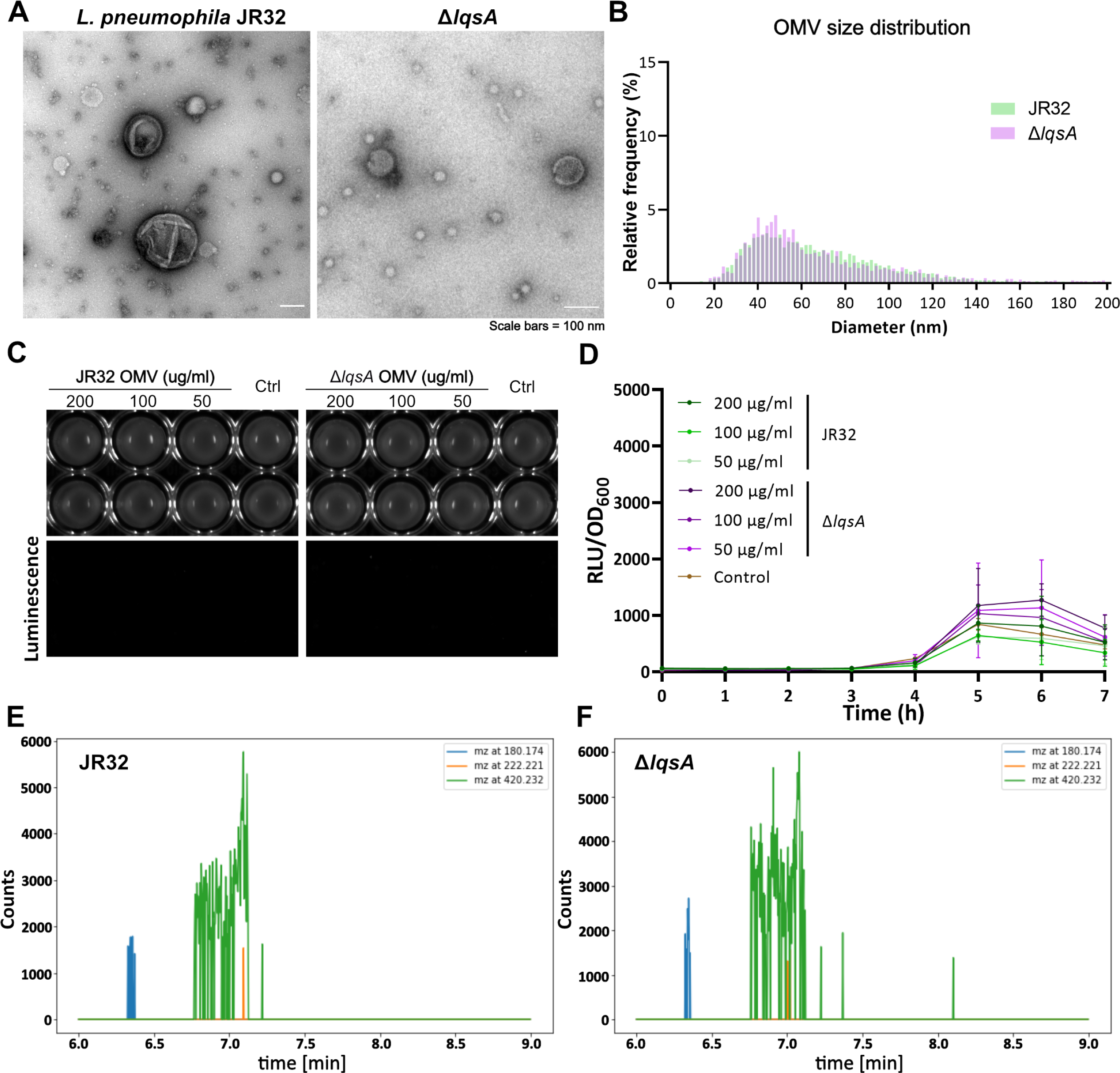
*L pneumophila* wild-type and Δ*lqsA* produce similar OMV populations. **(A)** Negative staining TEM images of OMVs derived from *L. pneumophila* JR32 (left) or Δ*lqsA* mutant strain (right). **(B)** Size distribution analysis of OMVs from *L. pneumophila* JR32 or Δ*lqsA*. **(C-D)** Luminescence of *V. cholerae* MM920 mixed with OMVs from *L. pneumophila* JR32 or Δ*lqsA* of the protein concentrations indicated (control: DMSO). Intensity was (**C**) visualized by a gel documentation system (15 min exposure time), or (**D**) measured by a plate reader (30°C, 7 h). RLU, relative light units. **(E-F**) LC-MS/MS analysis of OMVs of *L. pneumophila* JR32 or Δ*lqsA* (1 mg protein). Extracted ion chromatograms (EICs) of fragment ions at m/z 180.174 (light blue), 222.221 (orange), and 420.232 (green) did not indicate the presence of LAI-1 in these OMVs.

To increase the levels of LAI-1 produced by *L. pneumophila*, we sought to express *lqsA* from known strong promoters, which initially were tested as transcriptional fusions with *gfp*. Compared to the promoter P*_lqsA_*, the stationary phase promoters P*_flaA_* and P*_6SRNA_* indeed led to much higher levels of GFP (**Fig. 6A**). Moreover, judged from the production of GFP, P*_flaA_* and P*_6SRNA_* are strongly expressed in the early or later stationary growth phase, respectively, while P*_lqsA_*is barely expressed (**Fig. 6B**).

**Figure 6.**
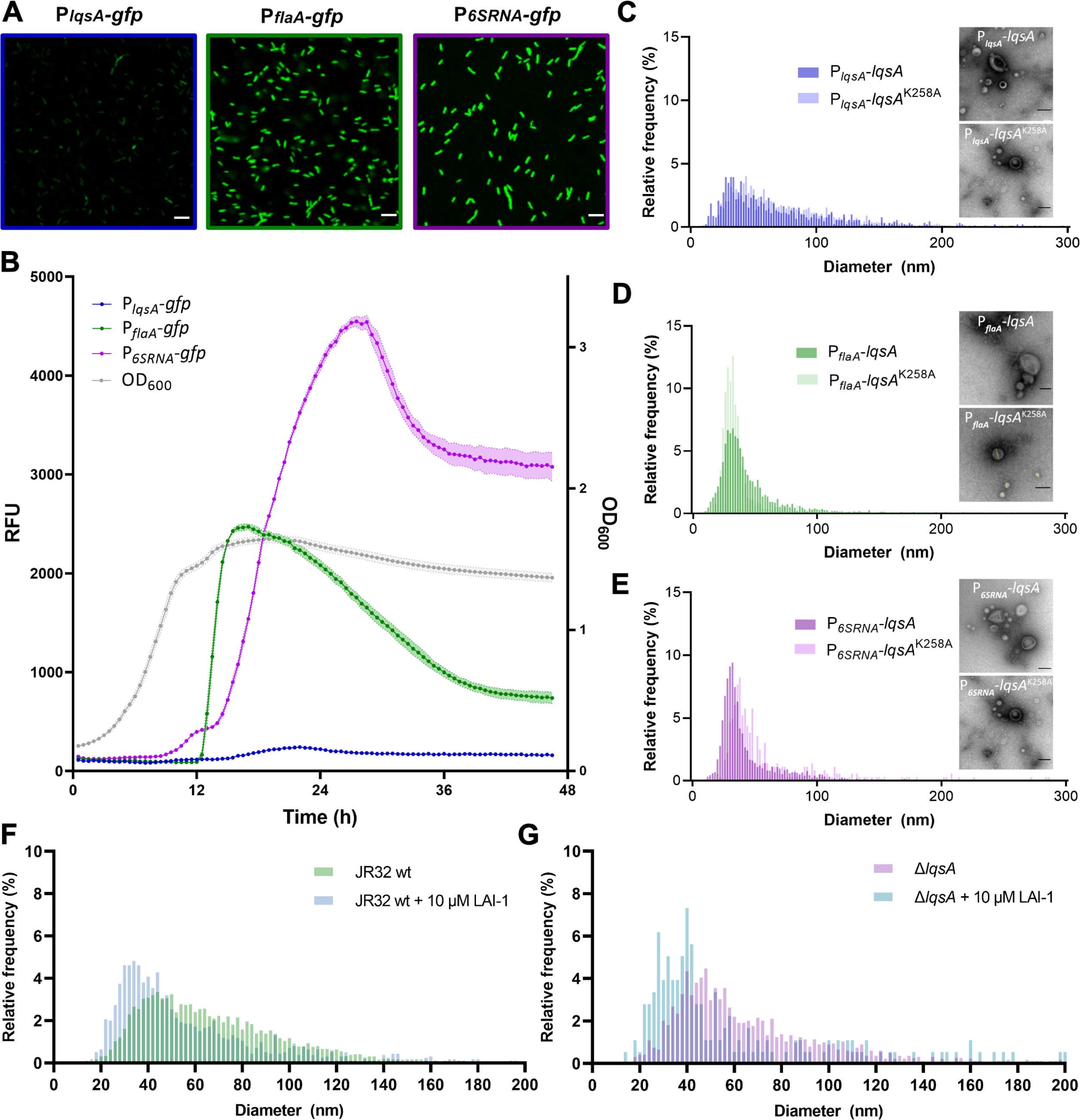
*LqsA* and LAI-1 reduce the median OMV size of *L. pneumophila*. GFP fluorescence intensity was measured by (**A**) confocal fluorescence microscopy of stationary phase *L. pneumophila* JR32 harboring pCM-5 (P*_lqsA_*-*gfp*), pCM-9 (P*_flaA_*-*gfp*) or pRM049 (P*_6SRNA_*-*gfp*), or (**B**) microtiter plate reader of stationary phase cultures diluted to an OD_600_ of 0.2 (RFU, relative fluorescence units; orbital shaking at 37°C, 48 h). The intensity of GFP fluorescence indicates that the promoter activity of P*_lqsA_* is considerably lower than that of P*_flaA_* or P*_6SRNA_*. **(C-E)** Size distribution and negative staining TEM of OMVs isolated from stationary phase cultures of *L. pneumophila* JR32 harboring (**C**) pAK014 (P*_lqsA_-lqsA*) or pTS-24 (P*_lqsA_-lqsA*^K258A^), (**D**) pMF04 (P*_flaA_-lqsA*) or pMF15 (P*_flaA_-lqsA*^K258A^), or (**E**) pMF03 (P*_6SRNA_-lqsA*) or pMF16 (P*_6SRNA_-lqsA*^K258A^). Strong overexpression of *lqsA* reduces the median OMV size. (**F**, **G**) 10 µM LAI-1 was added to (**F**) *L. pneumophila* JR32 or (**G**) Δ*lqsA* growing in mid-logarithmic phase and the OMV population was assessed. LAI-1 reduces the median OMV size.

Given the features of these promoters, we expressed *lqsA* under the control of P*_flaA_* or P*_6SRNA_* and compared the constructs to *lqsA* expressed under the control of its endogenous promoter P*_lqsA_*. The resulting *L. pneumophila* strains expressing *lqsA* (wild-type or a catalytically inactive mutant) under the control of P*_lqsA_*, P*_flaA_*, or P*_6SRNA_* showed the same morphology and grew indistinguishably in AYE broth (**Fig. S5**).

*L. pneumophila* expressing *lqsA* under the control of its endogenous promoter P*_lqsA_* (**Fig. 6C**) produced an OMV population similar to the corresponding strain overproducing a catalytically inactive LqsA mutant. The OMVs ranged in size from approximately 10 to 150 nm diameter. *L. pneumophila* expressing *lqsA* under the control of P*_flaA_* (**Fig. 6D**) or P*_6SRNA_*(**Fig. 6E**) produced OMV populations with individual OMV ranging in size from approximately 10 to 100 nm in diameter. The median OMV size was similar in strains expressing *lqsA* or *lqsA*^K258A^ from the same promoter: 67 nm/62 nm (P*_lqsA_*), and 37 nm/32 nm (P*_flaA_*). However, the median size of OMVs produced by *L. pneumophila* expressing *lqsA* under the control of P*_6SRNA_* was significantly smaller than the control (34 nm/45 nm). Overall, the OMVs produced by *L. pneumophila* expressing *lqsA* from its endogenous promoter P*_lqsA_* were bigger than the OMVs from the strains overproducing LqsA und control of P*_flaA_* or P*_6SRNA_*. This was observed independently of the catalytic activity of LqsA, and the reason is unknown.

Based on the finding that overexpression of *lqsA* under the control of P*_6SRNA_* results in smaller OMVs, we tested whether synthetic LAI-1 affects OMV formation. To this end, we added 10 µM LAI-1 to *L. pneumophila* JR32 or Δ*lqsA* growing in mid-logarithmic phase. Under these conditions, the median size of OMVs from *L. pneumophila* treated with LAI-1 (**Fig. 6F**) was significantly smaller compared to OMVs from untreated bacteria (**Fig. 6G**): JR32 – 47 nm/62 nm, Δ*lqsA* – 42 nm/58 nm), and therefore, exogenously added synthetic LAI-1 reduced the median OMV size. Taken together, *L pneumophila* OMV production is affected by the overexpression of *lqsA* or by exogenously added synthetic LAI-1.

### LAI-1 partitions to OMVs of L. pneumophila overexpressing lqsA

To assess whether LAI-1 produced by *L. pneumophila* strains overexpressing *lqsA* partitions into OMVs, we exposed the *V. cholerae* reporter strain MM920 to the corresponding OMV preparations (**Fig. 7A**). OMV preparations (50-250 µg protein/ml) purified from *L. pneumophila* strains expressing *lqsA* under control of P*_flaA_* or P*_6SRNA_* yielded robust, dose-dependent luminescence signals with a detection limit of ca. 100 µg protein/ml (**Fig. 7B**). In contrast, *L. pneumophila* strains expressing *lqsA* under control of its endogenous promoter P*_lqsA_* did not elicit any luminescence from the reporter strain. Accordingly, the overexpression of *lqsA* under control of strong promoters produces LAI-1, which partitions to OMVs and activates quorum sensing (luminescence) in a *Vibrio* reporter strain.

**Figure 7.**
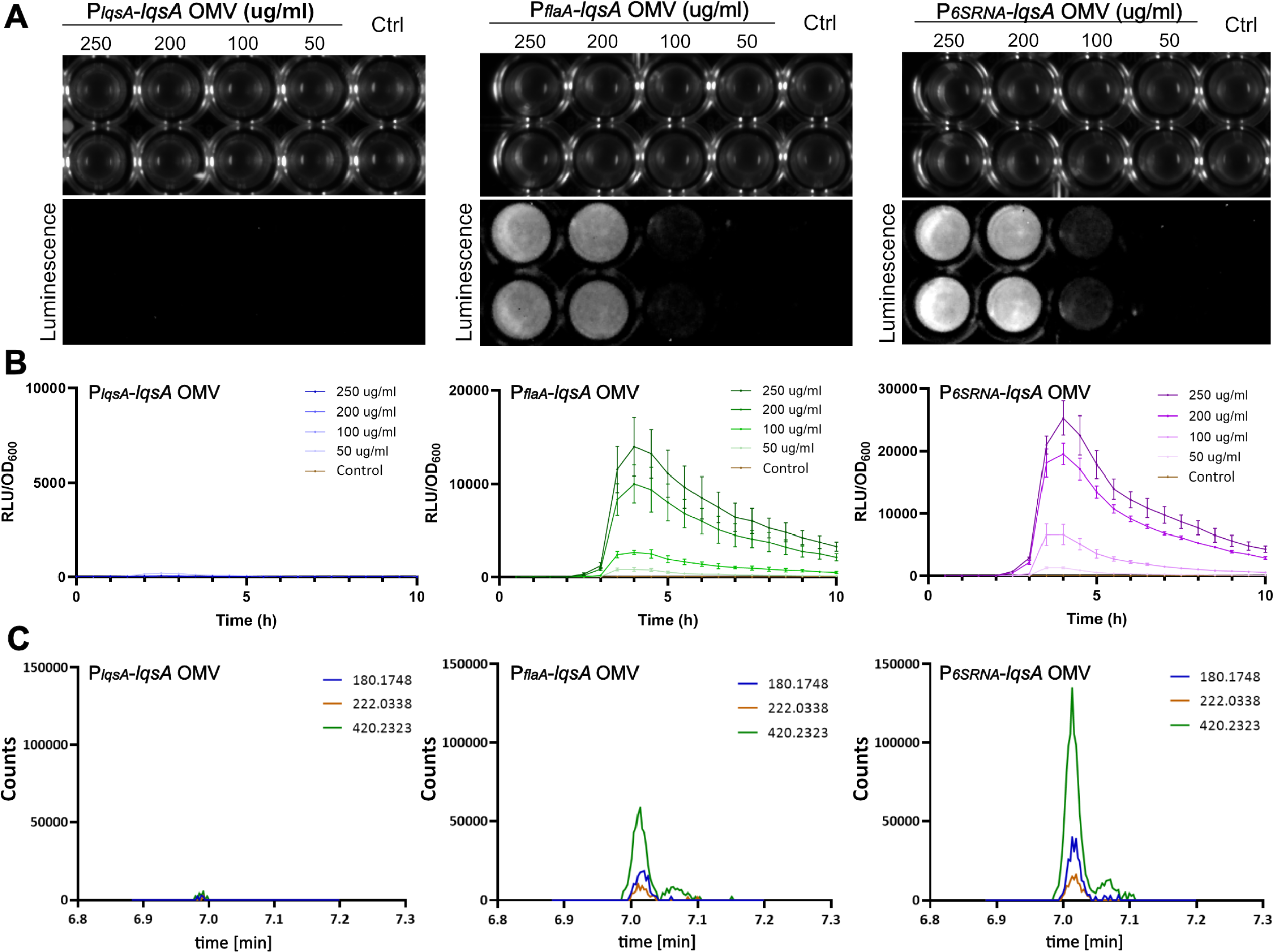
LAI-1 partitions to OMVs of *L. pneumophila* overexpressing *lqsA*. (**A**, **B**) Luminescence of *V. cholerae* MM920 mixed with OMVs from *L. pneumophila* harboring pAK014 (P*_lqsA_-lqsA*), pMF04 (P*_flaA_-lqsA*), or pMF03 (P*_6SRNA_-lqsA*) of the protein concentrations indicated (control: DMSO). Intensity was (**A**) visualized by a gel documentation system (15 min exposure time), or (**B**) measured by a plate reader (30°C, 10 h). RLU, relative light units. **(C)** LC-MS/MS analysis of OMVs from *L. pneumophila* harboring pAK014 (P*_lqsA_-lqsA*), pMF04 (P*_flaA_-lqsA*), or pMF03 (P*_6SRNA_-lqsA*) (1 mg protein). Extracted ion chromatograms (EICs) of fragment ions at m/z 180.174 (light blue), 222.221 (orange), and 420.232 (green) indicated a significantly higher amount of LAI-1 in OMVs of *L. pneumophila* harboring pMF04 (P*_flaA_-lqsA*) or pMF03 (P*_6SRNA_-lqsA*).

Analogous results were obtained upon detection of LAI-1 by LC-MS/MS (**Fig. 7C**, **Table 1**). LAI-1 was robustly detected and identified in OMV preparations from *L. pneumophila* expressing *lqsA* under control of P*_flaA_* or P*_6SRNA_*, but barely from an *L. pneumophila* strain expressing *lqsA* under control of P*_lqsA_*. Finally, the expression of a catalytically inactive *lqsA* mutant under control of any of these promoters did not yield MS-detectable LAI-1 in OMV preparations above background levels (**Fig. S6**, **Table 1**). In summary, the overproduction in *L. pneumophila* of wild-type but not catalytically inactive LqsA produces LAI-1, which partitions to OMVs.

### L. pneumophila OMVs containing LAI-1 inhibit the migration of D. discoideum

To address the question, whether *L. pneumophila* OMVs containing LAI-1 affect amoeba migration and promote inter-kingdom signaling, we treated *D. discoideum* with OMVs purified from *L. pneumophila* JR32 producing wild-type LqsA or the catalytically inactive mutant LqsA^K258A^ under control of P*_6SRNA_*(**Fig. 8**). Compared to the MB medium control (**Fig. 8A**), the migration of the amoeba was inhibited in a dose-dependent manner by OMVs purified from *L. pneumophila* expressing wild-type *lqsA* under the control of P*_6SRNA_* (**Fig. 8B**), but not by OMVs purified from *L. pneumophila* expressing catalytically inactive *lqsA* under the control of P*_6SRNA_* (**Fig. 8C**). At the highest OMV concentration used (1000 µg/ml protein), the velocity of the amoeba was reduced ca. 2-fold compared to the controls (**Fig. 8D**).

**Figure 8.**
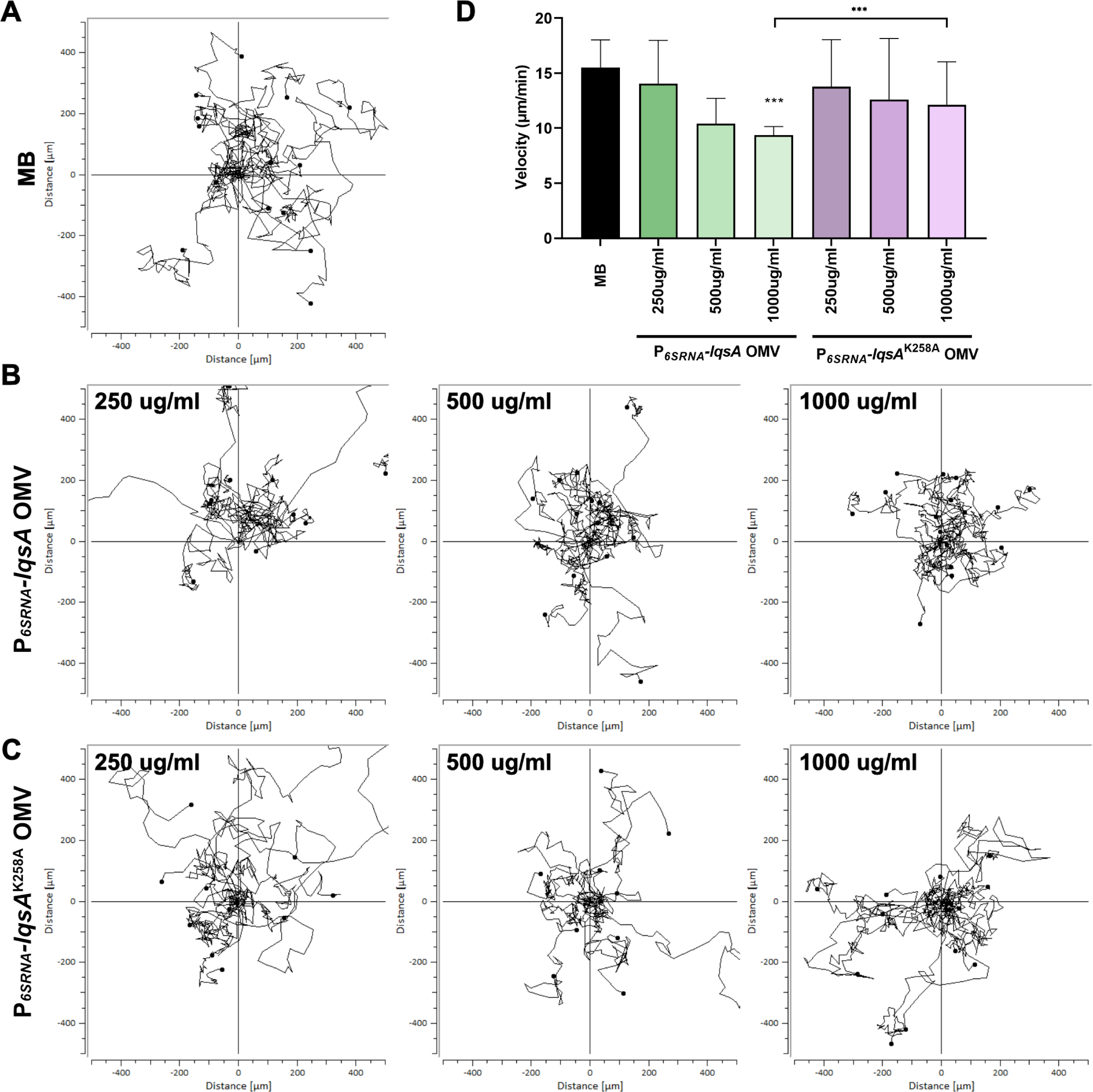
*L. pneumophila* OMVs harboring LAI-1 inhibit the migration of *Dictyostelium*. *D. discoideum* amoeba were treated with OMVs of *L. pneumophila* JR32 harboring pMF03 (P*_6SRNA_-lqsA*) or pMF16 (P*_6SRNA_-lqsA*^K258A^) with a final protein concentration of 250 µg/ml, 500 µg/ml, or 1000 µg/ml. For each sample, the migration of 10-15 amoeba was tracked over 2 h and the velocity was quantified. Migration trajectories of amoeba treated with (**A**) MB medium (control), (**B**) OMVs of *L. pneumophila* JR32 harboring pMF03 (P*_6SRNA_-lqsA*), or (**C**) pMF16 (P*_6SRNA_-lqsA*^K258A^). (**D**) Median of amoeba migration velocity. The data shown are velocity median of 10-15 amoeba per sample (***, *p* ≤ 0.001) and representative of 2 independent experiments.

### LqsA overexpression promotes intracellular growth of L. pneumophila

To test the hypothesis that LAI-1 produced by *L. pneumophila* might affect intracellular pathogen-host interactions, we analyzed the replication in macrophages of *L. pneumophila* strains constitutively producing GFP and expressing *lqsA* wild-type or catalytically inactive mutant under the control of P*_lqsA_*, P*_flaA_* or P*_6SRNA_*. These strains produced GFP at a comparable level (**Fig. S7**). Using these plasmids, LAI-1 was detected by the *V. cholerae* reporter strain MM920 in OMVs purified from strains expressing *lqsA* under the control of P*_flaA_* or P*_6SRNA_*, but not under the control of P*_lqsA_* (**Fig. 9A**). Intriguingly, upon infection of RAW 264.7 macrophages, the *L. pneumophila* strains overexpressing *lqsA* under the control of the P*_flaA_* or P*_6SRNA_* promoters replicated more efficiently in the host cells (**Fig. 9B**). Contrarily, the *L. pneumophila* strain expressing *lqsA* under control of it endogenous P*_lqsA_*promoter or strains expressing a catalytically inactive *lqsA* mutant did not show enhanced intracellular growth. Hence, the overexpression of *lqsA* but not a catalytically inactive mutant promotes intracellular replication of *L. pneumophila* in macrophages, indicating that intracellularly produced LA1-1 modulates the interaction in favour of the pathogen.

**Figure 9.**
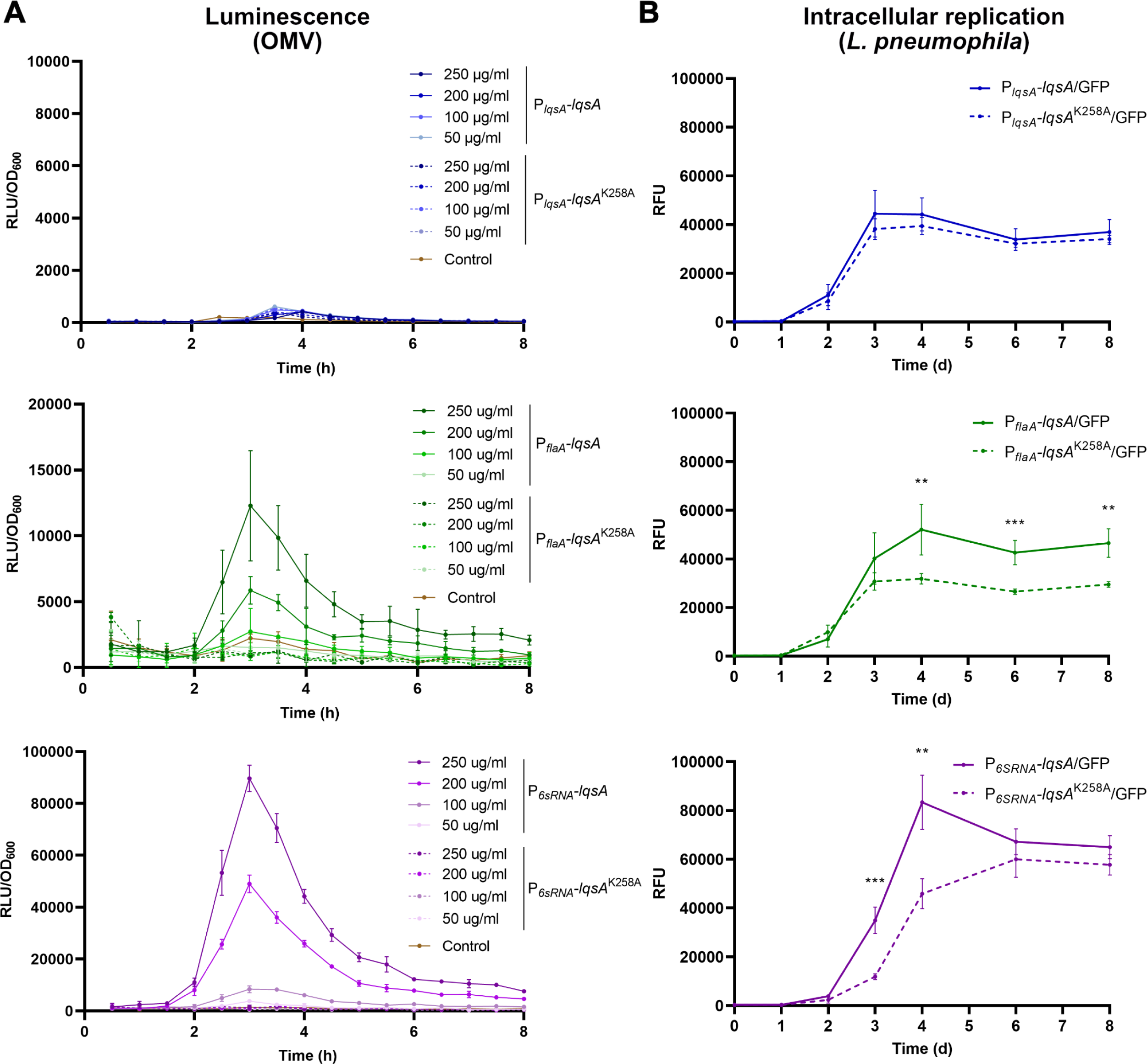
Overexpression of *lqsA* promotes intracellular replication of *L. pneumophila* in macrophages. (**A-C**) OMVs of GFP-producing *L. pneumophila* JR32 harboring (**A**) pMF21 (P*_lqsA_-lqsA*) or pMF22 (P*_lqsA_-lqsA*^K258A^), (**B**) pMF19 (P*_flaA_-lqsA*) or pMF20 (P*_flaA_-lqsA*^K258A^), or (**C**) pMF17 (P*_6SRNA_-lqsA*) or pMF18 (P*_6SRNA_-lqsA*^K258A^) were isolated, and LAI-1 was quantified using the *V. cholerae* reporter strain MM920 (OMV protein concentrations are indicated). (**D-F**) RAW 264.7 macrophages were infected (MOI 5) with GFP-producing *L. pneumophila* JR32 harboring (**D**) pMF21 (P*_lqsA_-lqsA*) or pMF22 (P*_lqsA_-lqsA*^K258A^), (**E**) pMF19 (P*_flaA_-lqsA*) or pMF20 (P*_flaA_-lqsA*^K258A^), or (**F**) pMF17 (P*_6SRNA_-lqsA*) or pMF18 (P*_6SRNA_-lqsA*^K258A^), and intracellular bacterial replication was followed by measuring GFP fluorescence over 8 d (RFU, relative fluorescence units). Data shown are means and standard deviations of triplicates (**, *p* ≤ 0.01; ***, *p* ≤ 0.001) and representative of 3 independent experiments.

## Discussion

In this study, we analyzed the mechanism of LAI-1 release as well as signaling effects of OMVs produced by *E. coli* and *L. pneumophila* strains expressing the autoinducer synthase gene *lqsA*. We found that the expression of *lqsA* in *E. coli* (**Fig. 3**) and *L. pneumophila* (**Fig. 5-7**) produces LAI-1, which partitions to OMVs. The OMV-localizing LAI-1 is bioactive in inter-bacterial as well as interkingdom signaling. Accordingly, LAI-1 in OMVs produced by *E. coli* triggers luminescence of *V. cholerae* MM920 (**Fig. 3CD**), induces expression of P*_flaA_*-*gfp* in *L. pneumophila* (**Fig. 3G**), and inhibits migration of *D. discoideum* (**Fig. 4**). Analogously, LAI-1 in OMVs produced by *L. pneumophila* triggers luminescence of *V. cholerae* MM920 (**Fig. 7**) and inhibits migration of *D. discoideum* (**Fig. 8**).

Gene expression analysis by qRT-PCR indicated that *lqsA* is expressed at the onset of the stationary growth phase (25). We observed a similar expression profile for *gfp* under the control of P*_lqsA_* (**Fig. S4**). However, compared to the strong stationary promoters P*_flaA_* and P*_6SRNA_*, the expression of P*_lqsA_* was very low (**Fig. 6**). Upon ectopic expression of *lqsA* under the control of different promoters, the strength of promoter activity correlated with the amount of LAI-1 detected by LC-MS/MS in OMVs: P*_6SRNA_* > P*_flaA_* > P*_lqsA_* (**Table 1**). No LAI-1 was detected in OMVs from *L. pneumophila* producing catalytically inactive LqsA, or in OMVs from *L. pneumophila* JR32 (**Table 1**). The expression of *lqsA* under control of the P*_tac_* promoter yielded ca. 30 times less LAI-1 in OMVs from *L. pneumophila* compared to *E. coli* (**Table 1**). Taken together, these findings correspond to the observation that *L. pneumophila* strains do not produce LAI-1 detectable by the α-hydroxyketone *V. cholerae* reporter strain MM920 or by LC-MS/MS, unless *lqsA* is overexpressed under control of P*_flaA_* or P*_6SRNA_* (**Figs. 2, 5**, **7**, **9**). Hence, under the conditions assessed, *L. pneumophila* tightly regulates the production of low amounts of LAI-1. It is not known whether *L. pneumophila* produces higher amounts of LAI-1 under different physiological conditions.

The results obtained here with OMV-localizing LAI-1 are similar to results obtained with synthetic LA1-1 for inter-bacterial signaling (33) and for inter-kingdom signaling (37). However, OMV-associated LAI-1 and synthetic LAI-1 might be solubilized in different micro-environments, i.e., in phospholipid bilayer vesicles or in micelles, respectively. Accordingly, the mode of action (membrane fusion vs. membrane insertion) and the bioavailability of OMV-associated LAI-1 and synthetic LAI-1 might differ substantially, and consequently, it is challenging to compare the efficacy of the two distinct LAI-1 delivery forms.

While we cannot exclude other transport mechanisms for LAI-1, overall, our findings provide ample evidence that LAI-1 is secreted by OMVs (**Fig. 1**, **3**, **7**). Since LAI-1 is rather hydrophobic, the compound likely partitions into the OMV membrane, but might also be present in the OMV lumen to some extent. As a general concept, packaging in OMVs of hydrophobic, low molecular weight signaling molecules solubilizes these compounds in aqueous environments and facilitates their distribution over rather large distances at biologically active concentrations (42). Hence, by being distributed through OMVs, LAI-1 and other hydrophobic bacterial signaling molecules experience an increased solubility and enhanced bioactivity.

The delivery through OMVs has been shown for a few hydrophobic quorum sensing compounds, which promote bacterial intra-species and inter-kingdom signaling (42). The opportunistic human pathogen *Pseudomonas aeruginosa* produces the compound 2-heptyl-3-hydroxy-4-quinolone (*Pseudomonas* quinolone signal, PQS), which is packaged into OMVs and mediates quorum sensing-dependent group activities in bacterial populations (52). Intriguingly, PQS stimulates its own OMV packaging and promotes OMV formation of *P. aeruginosa* (52). We obtained evidence that the overexpression of *lqsA* and synthetic LAI-1 decrease the median OMV size of *E. coli* (**Fig. 3**) and *L. pneumophila* (**Fig. 6**), and therefore, affect OMV formation. *Burkholderia thailandensis* hydroxyalkyl quinolone HMNQ is also released through OMVs (53) and *Vibrio harveyi* CAI-1 is packaged in OMVs in late stationary growth phase (54). CAI-1 in *V. harveyi* OMVs triggers quorum sensing phenotypes in CAI-1 non-producing *V. harveyi* and *V. cholerae*. Moreover, the marine coral pathogen *Vibrio shilonii* produces OMVs that contain quorum sensing molecules of the N-acylhomoserine lactone (AHL) family (55). Finally, *Paracoccus denitrificans* was shown to release the AHL signaling molecule N-hexadecanoyl-L-homoserine lactone (C16-HSL) *via* OMVs (56).

With regard to inter-kingdom signaling, LAI-1 is bioactive regardless of whether the compound is delivered extracellularly through purified OMVs from *E. coli* (**Fig. 4**) or *L. pneumophila* (**Fig. 8**), or whether the compound is produced intracellularly in host cells infected with *L. pneumophila* overexpressing *lqsA* (**Fig. 9**). These results are in agreement with the notion that the (unknown) eukaryotic LAI-1 receptor faces the extracellular milieu and – upon uptake of *L. pneumophila* – the lumen of the LCV. This topology would allow the recognition of LAI-1 upon extracellular delivery as well as upon intracellular production. Alternatively, the hydrophobicity of LAI-1 might allow for a fairly unrestricted membrane diffusion and receptor targeting.

The overexpression of *lqsA* but not a catalytically inactive mutant promotes intracellular replication of *L. pneumophila* in macrophages (**Fig. 9**). This result indicates that intracellular – and thus likely continuous or repeated – production of LA1-1 promotes pathogen-host interactions of *L. pneumophila*. Similarly, the overexpression of *lqsA* by intracellular *L. pneumophila* inhibited cell migration of host cells (37). While the inhibition of eukaryotic cell migration could be faithfully recapitulated by the addition of 10 µM synthetic LAI-1, synthetic LAI-1 added prior to or concomitantly with infected macrophages, did not enhance the uptake (37) or intracellular replication of *L. pneumophila*. Perhaps, LAI-1 needs to be produced in a continuous or repeated manner to exert intracellular effects. OMVs shed by intracellular *L. pneumophila* have been reported to inhibit fusion of phagosomes with lysosomes (51), and to promote bacterial replication in macrophages (47). These processes are mechanistically not well understood. Further studies will elucidate the pathways underlying the inter-kingdom detection of and response to LAI-1 by eukaryotic host cells.

### Experimental Procedures

#### Bacteria, cells, and reagents

The bacterial strains used are listed in **Table S1**. *E. coli* TOP10 was cultured overnight at 37°C in LB broth supplemented with 30 μg/ml Cam. *L. pneumophila* strains were grown for 3 days on charcoal yeast extract (CYE) agar plates (57), with or without chloramphenicol (Cam; 10 µg/ml) at 37°C. Bacterial colonies were used to inoculate liquid cultures (starting concentration OD_600_ of 0.1) in *N*-(2-acetamido)-2-aminoethanesulfonic acid (ACES)-buffered yeast extract (AYE) medium (58) at 37°C on a wheel (80 rpm) and grown for approximately 18 h, with Cam (5 µg/ml) added to maintain plasmids if required.

*V. cholerae* MM920 (22) was cultured overnight at 30°C in LB broth supplemented with 5 μg/ml tetracycline (Tet) prior to an experiment. *V. cholerae* MM920 lacks the autoinducer synthase gene *cqsA* (Δ*cqsA*) and the sensor kinase gene *luxQ* (Δ*luxQ*), and therefore, does not respond to the quorum sensing signal AI-2, and does produce but responds to the α-hydroxyketone compounds CAI-1 and LAI-1. Strain MM920 harbors plasmid pBB1, which contains the *luxCDABE* luciferase operon of *V. harveyi* and produces light upon detection of CAI-1 and LAI-1.

Murine macrophage-like RAW 264.7 cells (ATCC TIB-71, laboratory collection) were cultivated in RPMI 1640 medium (Life Technologies) supplemented with 10% heat-inactivated fetal calf serum (FCS; Life Technologies) and 1% glutamine (Life Technologies) at 37°C with 5% CO_2_ in a humidified atmosphere. The macrophages were grown in T75 flasks (Faust Laborbedarf AG) and split every second or third day.

*D. discoideum* strain Ax3 was cultivated in HL5 medium without glucose (Formedium) supplemented with D-maltose (Roth) at 23°C. The amoebae were grown in T75 flasks and split every third day.

(S)-LAI-1 was synthesized as described (37) and is referred to as “LAI-1” throughout the manuscript.

#### Molecular biology and plasmid construction

The plasmids utilized in this study are listed in **Table S1**. Cloning procedures followed standard protocols, with plasmid isolation performed using commercially available kits from Qiagen. DNA fragments were amplified using Phusion High Fidelity DNA polymerase, using the primers listed in **Table S2**. Gibson assembly was carried out using the NEBuilder HiFi DNA assembly kit (NEB). All constructed plasmids were validated through Sanger sequencing (Microsynth, Switzerland). To construct plasmid pAK14, P*_lqsA_*-*lqsA* was amplified by PCR using the LqsA-nat-fw and LqsA-mod-rev primers and genomic DNA as a template, and the fragment was cloned into plasmid pTS-28 cut with BamHI and MluI.

The plasmids pMF04 and pMF15 were created by replacing the *gfp* gene in pCM-9 (33) with either the *lqsA* gene or a point-mutated *lqsA*^K258A^ gene, resulting in the transcriptional fusions P*_flaA_*-*lqsA* and P*_flaA_*-*lqsA*^K258A^, respectively. Amplification of the *lqsA* or *lqsA*^K258A^ gene regions was performed using the oMF015 and oMF016 primers, with JR32 genomic DNA or pTS-24 as the respective template. The backbones for pMF04 and pMF15 were identical and obtained by amplifying pCM-9 using the oMF013 and oMF014 primers.

Similarly, plasmids pMF03 and pMF16 were constructed as transcriptional P*_6SRNA_*-*lqsA* and P*_6SRNA_*-*lqsA*^K258A^ fusions by replacing the *gfp* gene in pRH049 (36). The *lqsA* or *lqsA*^K258A^ gene regions were amplified using the oMF011 and oMF012 primers, with JR32 genomic DNA or pTS-24 as the respective template. The PCR products were then cloned into pCM-9, amplified by the oMF009/oMF010 primer set.

To construct plasmid pMF17-pMF21, plasmid pNT31 was used as the backbone by digesting with restriction enzyme BamHI. For pMF17 and pMF18, the insert DNA fragments were amplified with the oMF64 and oMF65 primers, using pMF03 and pMF16 as template. The insert DNA fragments of pMF19 and pMF20 were amplified with the oMF66 and oMF65 primers, using pMF04 and pMF15 as templates. The insert DNA fragment of pMF21 was amplified by oMF67 and oMF65 primers, using JR32 genomic DNA as template. To construct pMF22, plasmid pMF21 was used as the backbone by amplifying with the primers oMF70 and oMF71 primers. Its insert DNA fragment was amplified by oMF69 and oMF65 primers, using pMF16 as template. The amplified DNA insert fragments were cloned to their corresponding backbones using a NEBuilder HiFi DNA assembly kit.

#### Isolation of bacterial OMVs

To prepare the bacterial culture for OMV isolation, *L. pneumophila* strains were grown to late stationary phase (OD_600_ ∼5.0) in 420 ml AYE medium, supplemented with 5 µg/ml Cam if required. The *E. coli* strains harboring pTS-2 or pTS-10 were cultured in 420 ml of LB medium supplemented with 30 µg/ml Cam. The cultures were maintained at 37°C with shaking at 120 rpm for 16 h. To induce the expression of *lqsA* in *E. coli* harboring pTS-2 or pTS-10, 1mM IPTG was added to the bacterial culture, and the incubation was continued for another 5 h.

OMVs were isolated basically as described (54). A low-speed centrifugation step (1’500 × *g*, 15 min, 4°C) and subsequent membrane filtration (0.45 µm pore size, Huberlab) were carried out to remove bacterial cells and debris from the culture supernatant. The resulting filtrate was then divided equally into six 70 ml polycarbonate bottles (Beckmann Coulter, USA) and subjected to ultracentrifugation (150’000 × *g*, 2 h, 4°C) using an Optima L-100XP centrifuge with a 45 Ti fixed angle rotor (Beckmann Coulter, USA). The resulting pellets were washed with PBS buffer and subjected to another round of ultracentrifugation under the same conditions. This process yielded a clear yellowish solution containing the secreted OMVs. The pellets from the six polycarbonate bottles were resuspended in a total of 1 ml of PBS buffer, and the resuspended samples were filtered once again using 0.45 µm pore size syringe filters. The OMV yield was determined by quantifying their protein content using the Pierce™ BCA Protein Assay Kit (Thermo Scientific, USA) according to the manufacturer’s instructions. The OMV extracts were either used immediately for experiments or stored at −80°C for future use, with caution taken to avoid multiple freeze and thaw cycles.

#### Gene expression measurements with microplate reader

*L. pneumophila* strains harboring transcriptional *gfp* fusions were grown to late stationary phase (OD_600_ ∼5.0) in 3 ml AYE medium, supplemented with 5 µg/ml Cam if needed. Each bacterial culture was then diluted with fresh medium to OD_600_ ∼0.2, and 200 µl of diluted bacterial suspension was pipetted in triplicates in a Falcon 96-well tissue culture-treated microplate (Corning, USA). The growth curve (OD_600_) and GFP fluorescence intensity (excitation, 488 nm, emission, 528 nm; bottom read) was measured with a Biotek Cytation 5 microplate reader (Agilent Technologies, USA) every 0.5 h for 48 h at 37°C with continuous orbital shaking.

#### LC-MS/MS analysis

For the preparation of MS samples, OMVs extracted from different *L. pneumophila* or *E. coli* strains were subjected to the following procedure. Initially, an OMV sample was combined with dichloromethane in a glass tube at a volume ratio of 2:3. The mixture was then gently agitated on a bench rotator for 1 h at room temperature (RT), with a rotation speed of 4 rpm. Subsequently, the glass tube was allowed to stand undisturbed for 1 h, promoting the separation of the two liquid phases. The organic phase (lower phase) containing the target compounds was carefully collected. To remove the solvent, the collected organic phase was evaporated to dryness using a Concentrator Plus (mode “vacuum – high vapor”, V-HV) for 45 min at RT. The residual solution was reconstituted by adding 150 µl of acetonitrile. The reconstituted sample was then stored at −80°C until used for analysis.

Prior to analysis, samples were oximated using 0-pentafluorbenzyl hydroxylamine (O-PFB; Merck, Darmstadt, Germany). To this end a saturated solution of O-PFB in acetonitrile was prepared (ca 100 mg in 500 µl). 10 µl of derivatization reagent were added to 40 µl sample. Samples were incubated for 10 min at RT prior to analysis. LC separation was performed with a Thermo Ultimate 3000 UHPLC system (Thermo Scientific, CA) using a C_18_ reversed phase column (Kinetex XB-C18 column, particle size 1.7 µm, pore size 100 Å; dimensions 50 mm × 2.1 mm, Phenomenex, CA). solvent A was 0.1% (v/v) formic acid and solvent B was 0.1% formic acid in (acetonitrile:H_2_O, 95:5) at a flow rate of 400 µl min^-1^. Solvent B was varied as follows: 0 min, 40 %; 2 min, 40 %; 5 min, 100%; 7 min, 100%; 7.5 min, 40%; subsequently, the column was equilibrated for 2.5 min at the initial condition. Injection volume was 2 µl.

MS - Product Reaction Monitoring (PRM) analysis was carried out with a Thermo QExactice plus instrument (Thermo Fisher Scientific, Waltham, MA, USA) in the positive FTMS. Both experiments were performed with a mass resolution of 17’500 (m/z = 200). In case of PRM, a ramped collision energy (20, 25 and 30 eV) was applied. LA1-1 precursor ion at m/z 438.2 (unit resolution) applying high energy C-trap collision dissociation. A heated ESI probe was used applying following source parameters: vaporizer temperature, 380°C; sheath gas, 50 aux gas, 20; sheath gas, 50; sweep gas, 0; RF level, 50.0; capillary temperature, 275°C.

For reproducible estimation of absolute concentrations, external standards with concentrations 10, 100, 1000 and 8000 nM were measured with each batch. Standard curves were determined from summed peak areas of fragment ions at m/z 180.1748, 222.0338, and 420.2323 whereby fragment ion m/z 180.1748 was used as identification fragment. MS-level 1 analysis was used to confirm exact precursor ion mass.

#### Vibrio reporter assay

The *V. cholerae* strain MM920 was inoculated in LB liquid medium supplemented with 5 ug/mL Tet and incubated for 18 h at 37°C. The overnight culture was diluted with fresh medium to an OD_600_ of 0.25. The medium was supplemented with either synthetic LAI-1 (1-50 µM) or OMV samples (50-250 µg/ml protein concentration).

The mixtures were then transferred to a 96-well plate (Chemie Brunschwig AG, Switzerland) and bioluminescence (luminescence; bottom read) intensity was measured using a Biotek Cytation 5 microplate reader every 0.5 h for 8-10 h at 30°C with continuous orbital shaking. Images were captured after 4-5 h incubation (when bioluminescence intensity usually reached maximum levels) using the FluorChem^TM^ SP imaging system (Alpha-InnoTec, Switzerland) with an exposure time of 15 min. All experiments were performed in biological triplicates.

#### Single amoeba tracking

*D. discoideum* Ax 3 amoeba were seeded at a density of 2×10^4^ cells per well into an 8-well Ibidi chamber (Ibidi) and incubated for 3-4 h in HL5 medium to allow attachment to the bottom. The medium in each well was then replaced by a diluted OMV sample and incubated at 23°C for 1 h before microscopy imaging. *E. coli* OMVs were diluted using MB medium to a final protein concentration of 100 µg/ml, 250 µg/ml, or 500 µg/ml. *L. pneumophila* OMVs were diluted using MB medium to a final protein concentration of 250 µg/ml, 500 µg/ml or 1000 µg/ml.

#### Intracellular replication assay

For infection assays, RAW 264.7 macrophages were seeded at a density of 1×10^5^ cells per well onto 24-well tissue culture plates and incubated overnight in RPMI 1640/10% FCS medium. *L. pneumophila* was inoculated at an OD_600_ 0.2 in AYE medium and grown on a wheel at 37°C to stationary phase (OD_600_ ∼5.0, ∼2 × 10^9^ bacteria/ml). The bacterial cultures were then diluted to the desired density in pre-warmed RPMI 1640/10% FCS, and the infection was synchronized by centrifugation (1050 × *g*, 10 min, RT). After 1 h, the infected cells were washed 4 times with pre-warmed RPMI 1640/10% FCS, and further incubated for the time indicated. Depending on the experimental set-up, the infected cells were imaged live, lysed with 0.1% Triton X-100 and/or fixed with 4% paraformaldehyde (PFA).

#### Transmission electron microscopy

Transmission electron microscopy (TEM) was used for the morphological characterization and size distribution of OMVs extracted from different *L. pneumophila* or *E. coli* strains. TEM was carried out by using a FEI Tecnai G2 Spirit (FEI Company, Hillsboro, USA), operating at 120 Kv and equipped with Gatan Orius 1000 (4k × 2.6k pixels) camera (Gatan Inc., USA). FEI was used as the control software and Gatan Digital micrograph was applied for image acquisition.

Prior to TEM, negative staining was performed on OMV samples as follows: A droplet of 5 µl OMV sample (1000 µg/ml diluted with PBS buffer) was added onto copper a grid (300 square mesh, Formvar Carbon support film; Microtonano, Netherlands) and incubated for 60 sec to allow attachment. The liquid was gently blotted away using a Whatman filter paper. A drop of ddH_2_O was then added to wash the grid once. Then a drop of 1% uranyl acetate solution was added onto the grid and incubated for 45 sec. After blotting off the liquid, the grid was placed on a filter paper to let dry completely and transferred for TEM.

The size of the OMVs was manually measured from the images acquired by TEM using ImageJ. In each sample, a minimum of 1000 OMVs were individually measured for diameter, and the data was then used to plot the diameter distribution.

#### Confocal laser scanning microscopy

Fluorescent-based imaging was conducted with a Leica SP8 confocal laser scanning microscope (CLSM). The following imaging acquisition was used for all imaging experiments: white light laser at 488 nm (2% intensity), fluorescent signal was detected by a power HyD detector (emission range 500-520 nm), transmissive light was detected by a PMT detector with a gain of 380. For single amoeba tracking, three fields of interest were randomly selected for each sample and recorded continuously for 2 h with 2 mins time interval. Image analyses were performed using ImageJ and Chemotaxis and Migration Tool version 2.0 (Ibidi).

#### Statistical analysis

All statistical analyses were performed using GraphPad Prism version 7.01 for Windows, GraphPad Software, La Jolla California, USA (www.graphpad.com). Two-tailed student’s t tests were used to compare means between experimental samples and controls. For experiments with multiple samples, one-way ANOVAs were used with Tukey’s or Dunnett’s Multiple Comparison Test post hoc comparisons of means.

##### Abbreviations

Icm/Dot: intracellular multiplication/defective organelle trafficking
LAI-1: *Legionella* autoinducer-1
LCV: *Legionella*-containing vacuole
Lqs: *Legionella* quorum sensing
LvbR: *Legionella* virulence and biofilm regulator
GFP: green fluorescent protein
OMVs: outer membrane vesicles
TEM: transmission electron microscopy
T4SS: type IV secretion system

## Acknowledgements

We would like to thank Aline Kessler for cloning pAK14. This work was supported by the Swiss National Science Foundation (SNF; 31003A_175557, 310030_200706) to HH. Work in the groups of C.H. and J.S was supported by the Swedish research council (VR) grant 2019-05384 and the Deutsche Forschungsgemeinschaft (DFG) RTG 2581 (project number 417857878), respectively. The authors have no conflict of interest to declare.

---

**Table S1.**
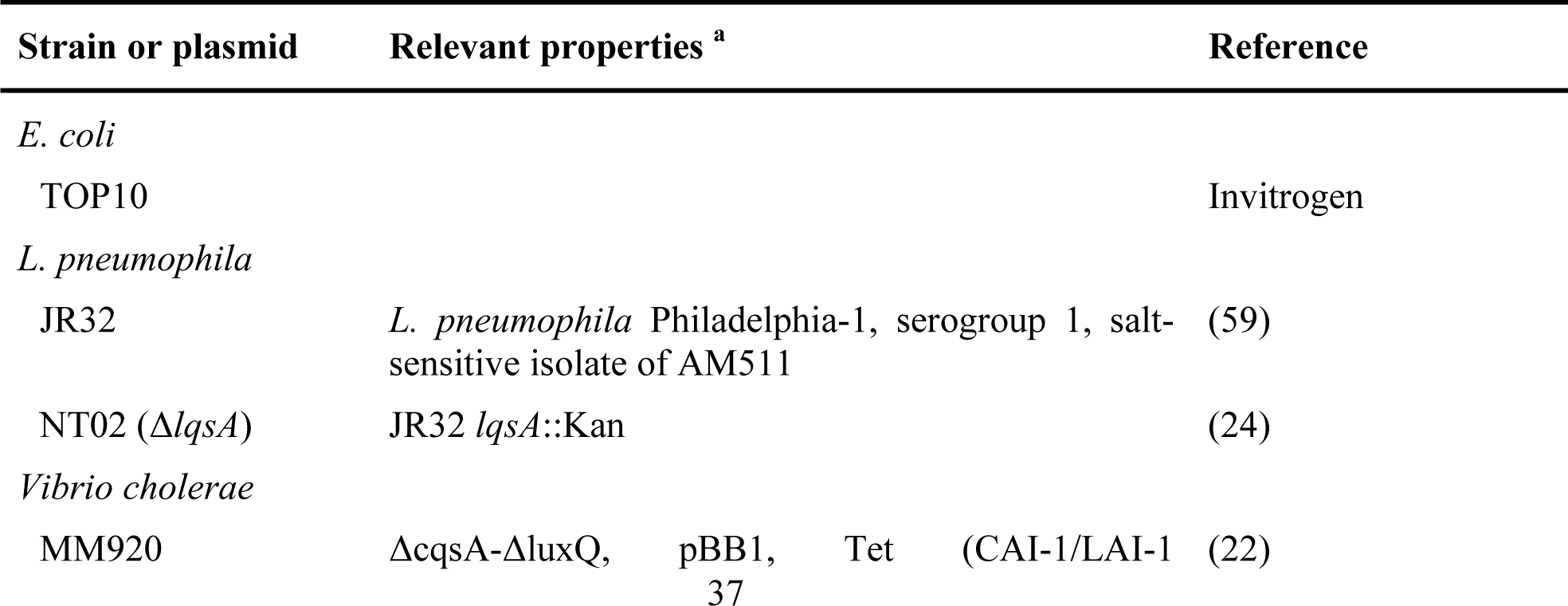
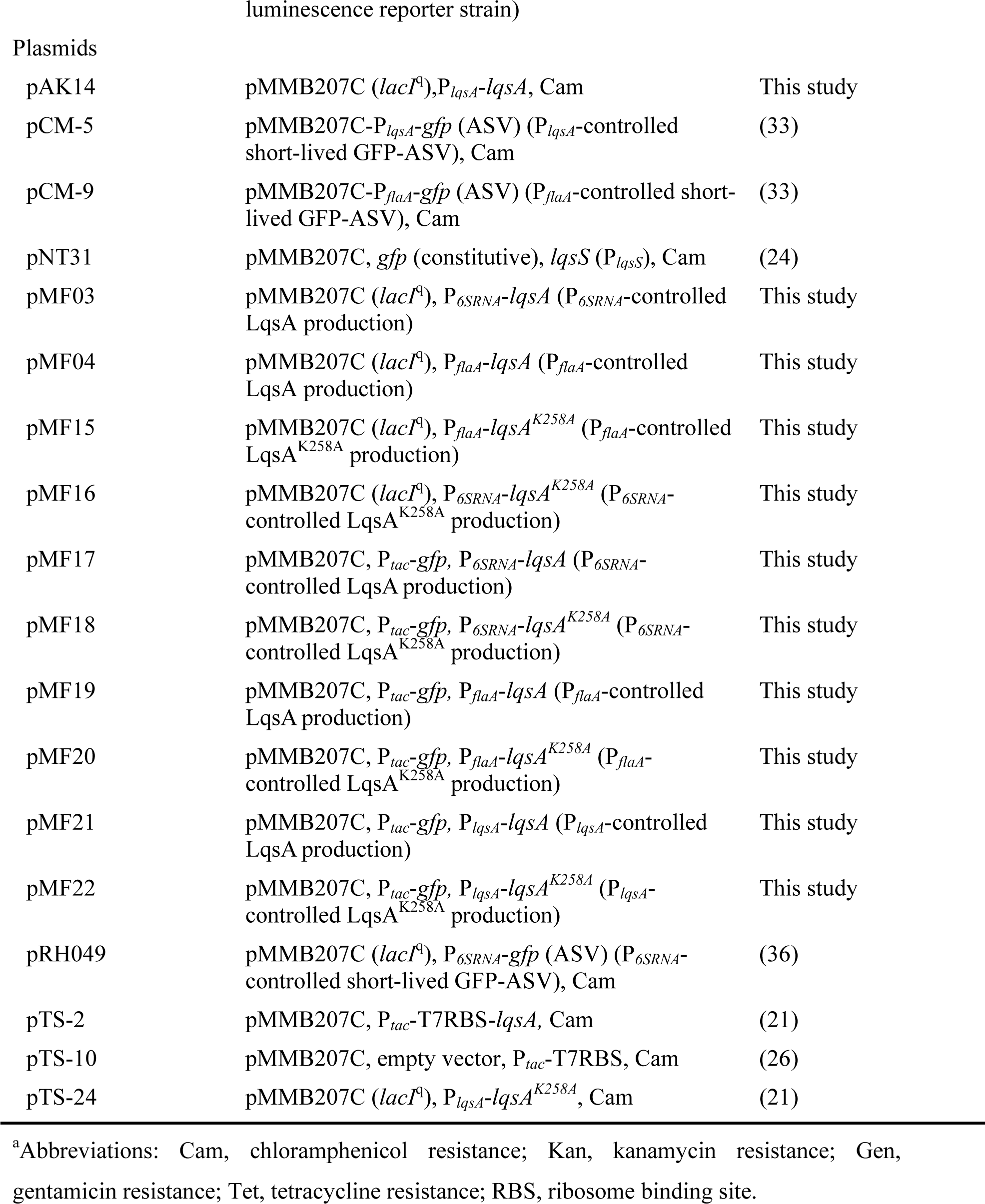
Bacterial strains and plasmids used in this study.

**Table S2.**
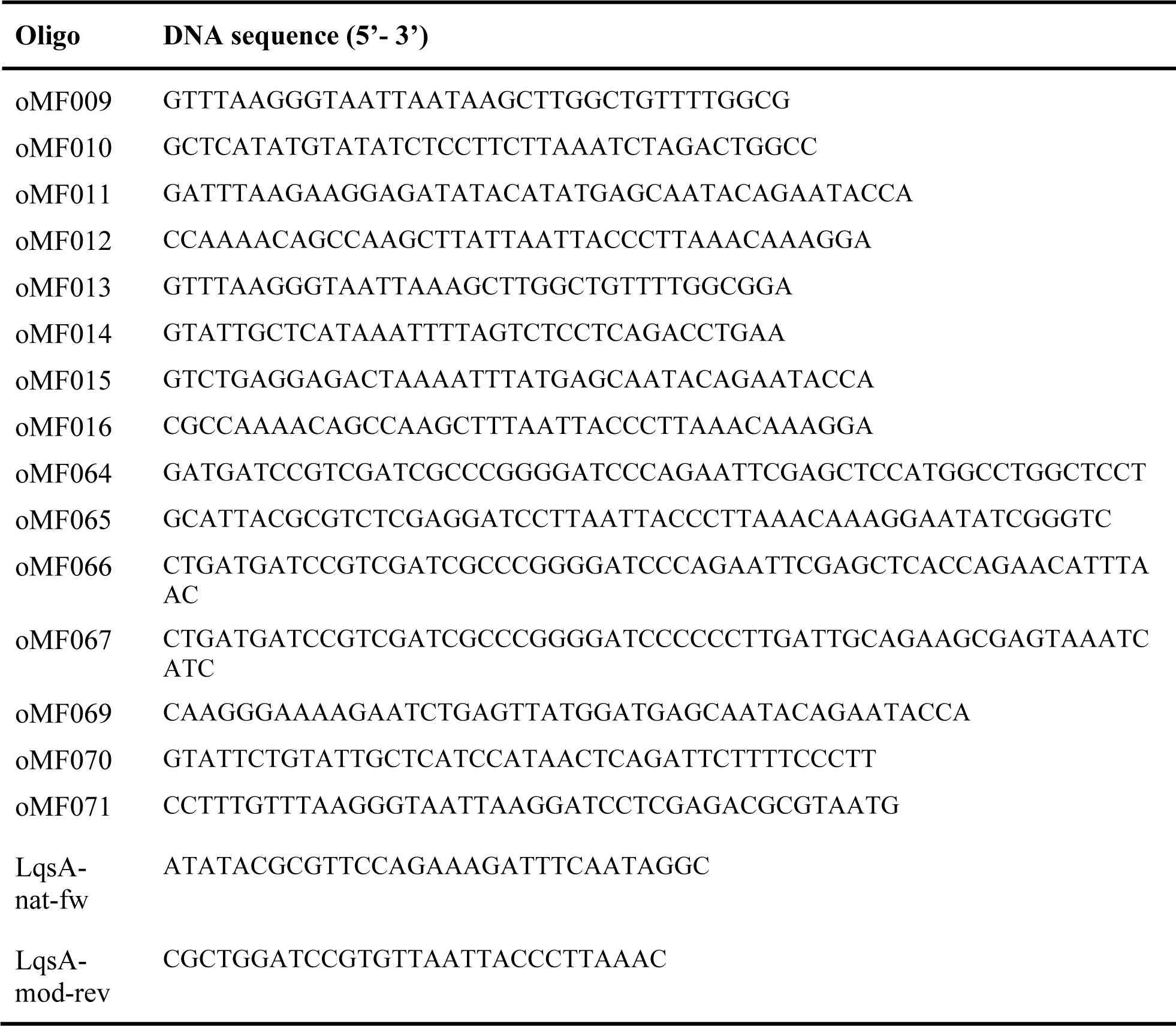
Primers used in this study.

**Figure S1. Detection of α-hydroxyketone signal in *L. pneumophila* supernatants by *V. cholerae* reporter strain MM920.** *L. pneumophila* harboring pTS-2 (P*_tac_*-*lqsA*) was grown in AYE broth for 14-19 h, and at the time points indicated, α-hydroxyketone activity in the supernatant was assayed by luminescence using the *V. cholerae* reporter strain MM920. The data are means and standard deviations of triplicates.

**Figure S2. MS fragmentation tree of oximated LAI-1.** The complete fragmentation tree of oximated LAI-1 (*Legionella* autoinducer-1, 3-hydroxypentadecane-4-one) is shown, and key fragments are highlighted with a blue star, or green, orange or blue triangles, respectively.

**Figure S3. Synthetic LAI-1 inhibits migration of *D. discoideum.*** *D. discoideum* amoeba were treated with 10 µM synthetic LAI-1. For each sample, the migration of 10-15 amoeba was tracked over 2 h and the velocity was quantified. Migration trajectories of amoeba treated with (**A**) MB medium (control), orn (**B**) 10 µM synthetic LAI-1. (**C**) Median of amoeba migration velocity. The data shown are velocity medians of 10-15 amoeba per sample (*, *p* ≤ 0.02) and representative of 3 independent experiments.

**Figure S4. Correlation of *L. pneumophila* growth and *lqsA* expression.** (**A**) Stationary phase *L. pneumophila* JR32 harboring pCM-5 (P*_lqsA_*-*gfp*) was diluted to an OD_600_ of 0.2 and grown in AYE medium in a 96-well plate while measuring OD_600_ and GFP fluorescence. OMVs were isolated from stationary phase *L. pneumophila*. (**B**) The growth of JR32 harboring pCM-5 (P*_lqsA_*-*gfp*) was visualized using confocal microscopy, with each image captured at one-hour intervals to monitor the progressive changes over time.

**Figure S5. Overexpression *lqsA* does not affect *L. pneumophila* growth in broth.** GFP-producing *L. pneumophila* JR32 harboring (**A**) pMF21 (P*_lqsA_-lqsA*) or pMF22 (P*_lqsA_- lqsA*^K258A^), (**B**) pMF19 (P*_flaA_-lqsA*) or pMF20 (P*_flaA_-lqsA*^K258A^), or (**C**) pMF17 (P*_6SRNA_-lqsA*) or pMF18 (P*_6SRNA_-lqsA*^K258A^) were grown in AYE medium (37°C, OD_600_).

**Figure S6. LAI-1 analysis by *V. cholerae* reporter strain of OMVs from *L. pneumophila* expressing catalytically inactive *lqsA*.** *V. cholerae* MM920 was treated with OMVs (protein concentrations indicated) from *L. pneumophila* harboring pTS-24 (P*_lqsA_-lqsA*^K258A^), pMF15 (P*_flaA_-lqsA*^K258A^), or pMF16 (P*_6SRNA_-lqsA*^K258A^), and luminescence was measured by a plate reader (30°C, 10 h). RLU, relative light units.

**Figure S7. GFP fluorescence intensity of *L. pneumophila* strains overexpressing *lqsA*.** GFP-producing *L. pneumophila* JR32 harboring (**A**) pMF21 (P*_lqsA_-lqsA*) or pMF22 (P*_lqsA_- lqsA*^K258A^), (**B**) pMF19 (P*_flaA_-lqsA*) or pMF20 (P*_flaA_-lqsA*^K258A^), or (**C**) pMF17 (P*_6SRNA_-lqsA*) or pMF18 (P*_6SRNA_-lqsA*^K258A^) were grown in AYE medium for 21 h to stationary phase, and fluorescence micrographs were taken by confocal microscopy.

